# Deep Learning Improves Macromolecule Identification in 3D Cellular Cryo-Electron Tomograms

**DOI:** 10.1101/2020.04.15.042747

**Authors:** E. Moebel, A. Martinez-Sanchez, L. Lamm, R. Righetto, W. Wietrzynski, S. Albert, D. Larivière, E. Fourmentin, S. Pfeffer, J. Ortiz, W. Baumeister, T. Peng, B.D. Engel, C. Kervrann

## Abstract

Cryo-electron tomography (cryo-ET) visualizes the 3D spatial distribution of macromolecules at nanometer resolution inside native cells. While this label-free cryogenic imaging technology produces data containing rich structural information, automated identification of macromolecules inside cellular tomograms is challenged by noise and reconstruction artifacts, as well as the presence of many molecular species in the crowded volumes. Here, we present a computational procedure that uses artificial neural networks to simultaneously localize with a multi-class strategy several macromolecular species in cellular cryo-electron tomograms. Once trained, the inference stage of DeepFinder is significantly faster than template matching, and performs better than other competitive deep learning methods at identifying macromolecules of various sizes in both synthetic and experimental datasets. On cellular cryo-ET data, DeepFinder localized membrane-bound and cytosolic ribosomes (~3.2 MDa), Rubisco (~540 kDa soluble complex), and photosystem II (~550 kDa membrane complex) with comparable accuracy to expert-supervised ground truth annotations. Furthermore, we show that DeepFinder is flexible and can be combined with template matching to localize the missing macromolecules not found by one or the other method. The DeepFinder algorithm is therefore very promising for the semi-automated analysis of a wide range of molecular targets in cellular tomograms, including macromolecules with weights of 500-600 kDa and membrane proteins.

## INTRODUCTION

Decades of research in cell biology have revealed that cellular processes are performed by groups of interacting macromolecules in a crowded environment. This is in contradiction with the previous cell models where macromolecules were considered as isolated objects floating randomly in the cytoplasm^1^. Deciphering the underlying interaction mechanisms is thus of paramount importance to gain a deeper understanding of the cell. Cryo-electron tomography (cryo-ET) can provide new insights into molecular interactions by producing 3D views of the native cellular interior at sufficient resolution to identify macromolecules. Unlike light microscopy, cryo-ET lacks specific markers, so the structures of the macromolecules themselves must be used for identification. However, cryo-ET has an advantage over light microscopy in that it visualizes everything in the cell, not just the tagged molecules of interest. With enough resolution and powerful computational tools for structural identification, cryo-ET has the potential to build a complete molecular atlas of the cell.

Cryo-ET data is generated by the following steps. First, the samples are vitrified in order to preserve both the native structures and spatial distribution of macromolecules in the cell. For most cells, a thinning step is required, which can be accomplished with focused ion beam milling^2^. Subsequently, the thin biological material is loaded into the transmission electron microscope for acquisition of a tilt-series, which serves as input to 3D reconstruction algorithms^3;4;5^. During tilt-series acquisition, the specimen is rotated around an axis perpendicular to the electron beam and imaged from multiple perspectives. The tilt-series must be acquired with low electron dose because frozen biological material is easily damaged by the electron beam. Unlike the heavy metal contrasting agents used in conventional electron microscopy, the organic molecules found in frozen cells have low contrast against the water background. Combined with the limited electron dose used for imaging, cryo-ET data has a very low signal-to-noise ratio. Furthermore, due the geometry of the EM grid and increasing thickness of the sample at high tilts, tilt-series are typically restricted to roughly ±60 degrees. As a result, the reconstructed tomograms suffer from a missing wedge of information in Fourier space. This missing wedge artifact causes anisotropic resolution in the 3D volumes, with delocalization of densities along the Z-direction, as well as loss of information in the X–Y plane along the direction perpendicular to the tilt axis^6;7^ The missing wedge and low signal-to-noise ratio, combined with the highly crowded environment of the cell, pose significant challenges to the identification of macromolecules in cellular tomograms.

One well-established method for localizing macromolecules in cryo-ET data is template matching^8^, where a low-resolution template depicting the macromolecule of interest is comprehensively scanned through the tomogram. High cross-correlation scores indicate potential particle positions, from which subvolumes are extracted for downstream averaging procedures. While template matching is relatively efficient for localizing large complexes such as ribosomes, it is necessary to apply a series of iterative searching, alignment, and classification steps to identify smaller molecules^9^. Additional difficulties arise when template matching is used to localize several macromolecular species that are structurally similar, or to differentiate specific states of the same macromolecular species (e.g., membranebound vs. cytosolic ribosomes). Template matching is applied to separately localize all macromolecules of a single species (mono-class procedure). Nevertheless, dedicated classification steps^10;11^ are required to differentiate true particles from false positives, as well as to subdivide these particles into structurally distinct sub-classes. Classification remains a challenging problem in cryo-ET if the number of considered classes or subclasses is high. Currently, complex and time-consuming processing chains are routinely used to localize macromolecules in cellular volumes, with a single template matching or subtomogram classification round typically taking 10-30 hours of computation on specialized CPU clusters. As a result, this whole procedure is most often used to analyze only a few classes or subclasses of particles in the same volume. However, cryo-electron tomograms contain many more macromolecular species embedded within the crowded cellular environment, hidden by noise and reconstruction artifacts. To address this challenging issue, powerful new pattern recognition techniques are required.

In this paper, we describe a deep learning-based framework to quickly identify macromolecules in cryo-electron tomograms. Deep learning^12^ is revolutionizing various fields of data processing, including computer vision^13^, language processing^14^, image classification^15^ and segmentation^16^, and object recognition^17^. In bioimage analysis, convolutional neural networks (CNN) have produced spectacular results^18;19;20;21;22^, including in super-resolution microscopy^23^ and in light microscopy image denoising^24^. Briefly, a CNN is defined as an architecture composed of successive connected neuron layers. The neurons are small processing units capable of performing linear and nonlinear operations. Each neuron is controlled by parameters that are optimized during the learning process. In the case of CNNs, the neurons are applied in a convolutive manner, which enables the network to deal with the information redundancy of neighboring pixels. Thus, a neuron can be thought of as a filter, and a neuron layer as a filter bank. Each layer automatically extracts various and specific features from the data. Applying the layers sequentially enables the network to progressively compute high-level features, which results in a hierarchical or multiscale representation of the data. In human face recognition, the first layers typically encode basic features such as image contours/edges and textures, which allows the next layers to gradually capture more complex shapes (e.g., eyes, ears), and object ensembles (e.g., faces). These multiscale representations are learned from the data, and can be generated faster than conventional handcrafted representations, which require time-consuming manual annotation by experts.

CNNs have recently been investigated for learning high-level generic features in electron microscopy. Several algorithms based on deep learning techniques have been developed for 2D particle picking in single-particle cryo-EM^25^, including DeepPicker^26^, AutoCryoPicker^27^, crYOLO^28^, Topaz^29^, and Warp^30^. In cellular cryo-ET, the algorithm proposed by Chen *et al*., was implemented for supervised 2D segmentation^31^, but it was not designed to handle complex environments (e.g., crowded cell) and requires an additional classification step to achieve satisfactory results. A promising 3D processing approach for the supervised classification of subtomograms has been proposed in Che *et al*. ^32^. However, this algorithm, which we have evaluated on our dataset, assumes that the molecular complexes of interest have already been found by another method. To overcome the limitations of the aforementioned approaches^26;31;32^, we propose a 3D deep learning method to identify macromolecules within a native crowded cellular environment as illustrated in Fig. 1B. Our DeepFinder algorithm can handle multiple macromolecular species in one pass, which we demonstrate actually improves the performance of CNNs in 3D cryo-ET, identifying small particles that template matching struggles to detect. Moreover, complex and time-consuming post-classification steps are no longer required to produce reliable results. We also show that DeepFinder is very flexible, and can be efficiently combined with template matching to improve localization sensitivity on crowded cellular cryo-tomograms.

**Figure 1:**
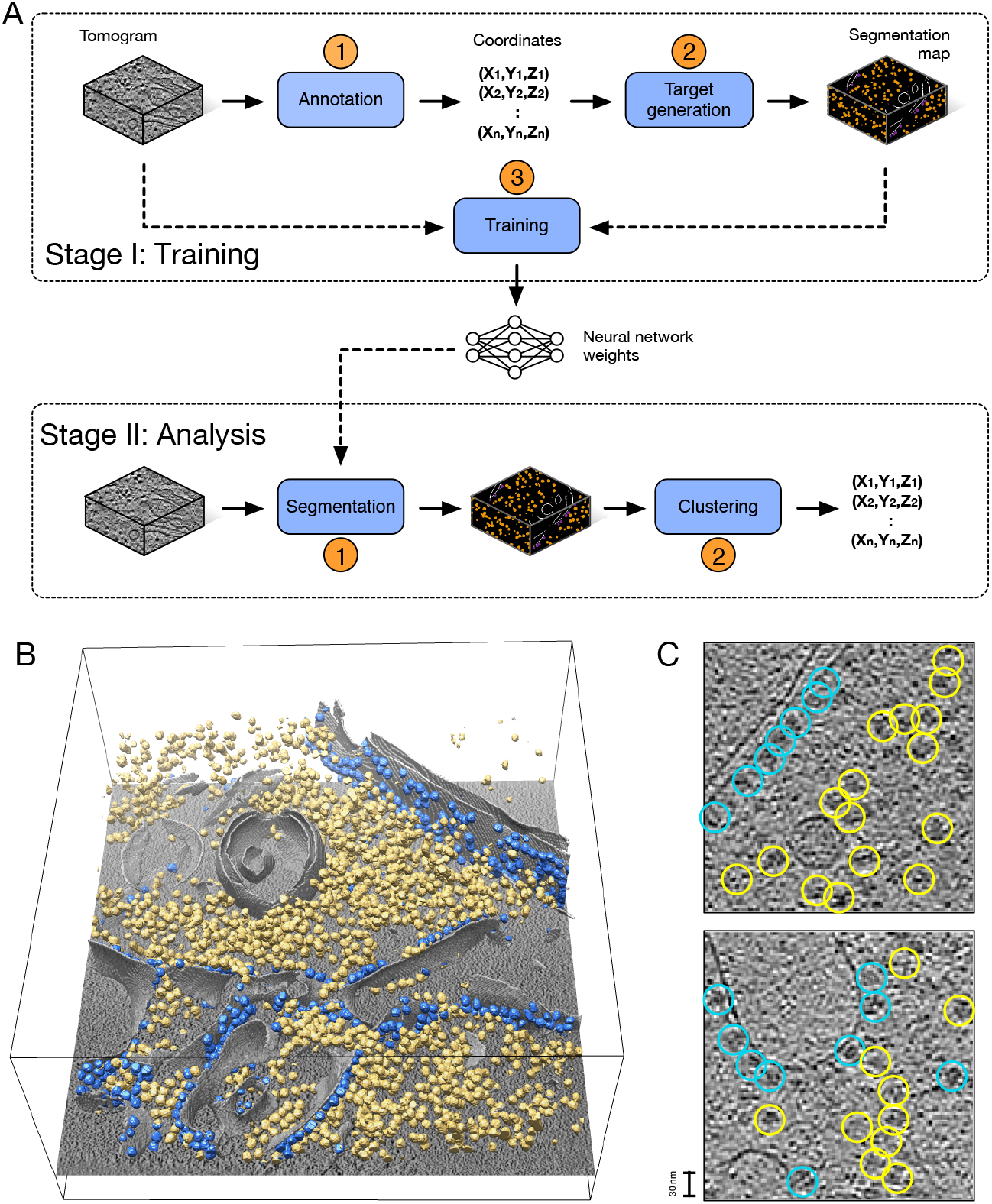
Overview of DeepFinder. (A) The DeepFinder workflow consists of training stage (Stage I) and an analysis (or inference) stage (Stage II). These two stages correspond to five steps (represented by blue boxes) to locate macromolecular complexes within crowded cells. The DeepFinder package provides a graphical interface for each step to to increase the accessibility of the algorithm. Meanwhile, each step can be executed with scripts using the API. (B) Ribosome localization with DeepFinder in a cryo-electron tomogram of a *Chlamydomonas reinhardtii* cell. Tomographic slice with superimposed segmented cell membrane (gray) and ribosomes classified with respect to their binding states: membrane-bound (blue) and cytosolic (yellow). (C) Tomographic slices showing coordinates of detected ribosomes (colors correspond to (B)). The positions and classes were determined by analyzing the segmentation map shown in (B).

## RESULTS

### Overview of our 3D deep learning-based approach

We propose DeepFinder, a flexible and powerful algorithm based on 3D convolutional neural networks that in one pass can robustly localize macromolecules of several different species, with various sizes and shapes, within cryotomograms. DeepFinder, which showed high performance on the synthetic SHREC’19 challenge dataset^33^ (https://www2.projects.science.uu.nl/shrec/cryo-et/2019/) and SHREC’20 challenge^34^ dataset, is an algorithm consisting of a training stage (Stage I) and an analysis (or inference) stage (Stage II) (see details in Methods and Fig. 1A). The analysis stage is a two-step procedure. In the first step, a supervised 3D CNN-based method is used to classify the 3D tomogram voxels into different categories or sub-categories of macromolecules. In the second step, a clustering algorithm is applied to aggregate voxels into groups and determine the location of particles (gravity center of clusters) in the volume (Fig. 1B). The particles are further picked and used for structure determination through subtomogram averaging. DeepFinder is a user-friendly algorithm implemented in Python (Fig. 1A) with a graphical user interface (see Supplementary Fig. 6 for illustration). The code can be downloaded for free from our GitLab website (https://gitlab.inria.fr/serpico/deep-finder) along with accompanying documentation (https://deepfinder.readthedocs.io/en/latest/). DeepFinder is embedded into the new release of Scipion (https://scipion-em.github.io/docs/linking_software/api/deepfinder/deepfinder.html), an open-source image processing framework for cryo-electron microscopy (http://scipion.i2pc.es/).

### DeepFinder identifies multiple macromolecular species at once

DeepFinder is built upon a multi-class network (see Methods and Supplementary Fig. 7). The underlying architecture is based on a U-net architecture^16;20^, as illustrated in Supplementary Fig. 8. We found that a multi-class strategy is more beneficial than identifying macromolecule classes separately. To demonstrate this point, we evaluated the performance of our multi-class network versus 12 mono-class versions of our network on the SHREC’19 dataset (Dataset #1).The SHREC’19 dataset is composed of 10 synthetic tomograms, each containing 12 different classes of macromolecules, with different sizes and structures^33^. These macromolecules have been categorized by size into 4 groups (large, medium, small, tiny) by the organizers. The number of macromolecules per class and per tomogram varies slightly. On average, a tomogram contains 208 macromolecules per class. As shown in Fig. 2B, the binary networks have a similar performance when compared to multi-class networks for larger macromolecules (e.g., macromolecules with the following Protein Data Bank (PDB) entry identifiers: 4b4t, 4d8q), but perform worse (or fail) for the smaller ones (e.g., 1s3x, 3gl1, 3h84 and 3qm1). Overall, the multi-class training improves classification accuracy over binary classification. These results also suggest that smaller macromolecules and under-represented classes can benefit from features learned from classes of larger macromolecules.

**Figure 2:**
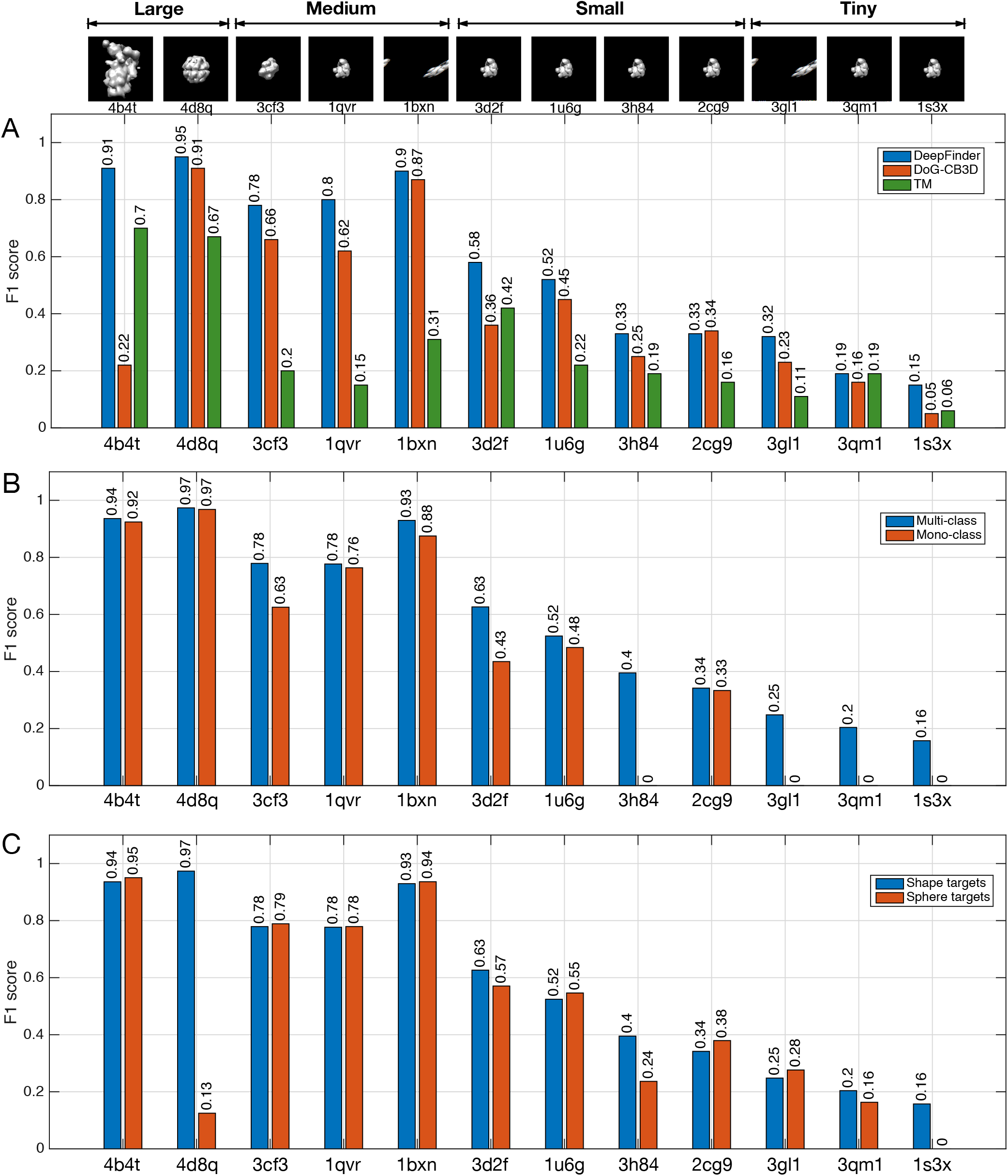
Analysis of algorithm performance on the synthetic dataset (SHREC’19 challenge). (A) Performance (*F*_1_-score) of DeepFinder, DoG-CB3D and template matching algorithms and ability to discriminate between 12 classes/subclasses of macromolecules. The tiny macromolecules are 1s3x (human Hsp70 ATPase domain), 3qm1 (Lactobacillus Johnsoni Cinnsmoyl esterase LJ0536 S106A mutant in complex Ethylferulate Form II), and 3gl1 (Saccharomyces cerevisiae ATPase domain of Ssb1 chaperone). The scores of template matching were provided by the SHREC’19 challenge organizers (Utrecht University, Department of Information and Computing Sciences and Department of Chemistry). (B) Performance of DeepFinder implemented as a multi-class network architecture and as an architecture made of 12 binary networks. These two architectures differ only by the number of output neurons. (C) Influence of the training target generation method (“shapes” versus “spheres”). In the case of “shapes”, the exact shapes of the macromolecules have been used to annotate the tomograms. In the case of spheres”, the shape and the orientation of macromolecules are not needed to generate the training targets.

In the SHREC’19 challenge, seven different methods (including DeepFinder) were compared to evaluate the performance of deep learning techniques on challenging synthetic 3D cryo-electron tomograms. Participants had access to 9 out of 10 ground truth tomograms, and then the algorithms were tested on the final unseen tomogram. DeepFinder achieved the best global *F*_1_-score (the highest possible value of an *F*_1_-score is 1.0 and the lowest possible value is 0) (see evaluation in Methods) among all competing methods^33^. The most serious competitor to DeepFinder was the DoG-CB3D method^32^, which separates the localization and classification tasks. In Fig. 2A, we report the results obtained on the SHREC’19 dataset with DeepFinder and DoG-CB3D. Unlike DoG-CB3D, DeepFinder outperformed template matching for all macromolecular species in the benchmark dataset. The template matching results were obtained with the PyTom software^35^ described in Appendix A. Performance is correlated with macromolecular size (Fig. 2A). This is especially true for template matching, where the smallest complexes had *F*_1_-scores close to 0. Even so, template matching is still a competitive method for detecting the largest macromolecules. All these results were confirmed on the SHREC’20 dataset^34^. In this most recent challenge, more attention was paid to the image formation process. In particular the contrast transfer function (CTF) was carefully designed in order to generate more realistic tomograms. Meanwhile, the organizers provided the results obtained with template matching described in Appendix A. In 2020, DeepFinder was ranked second, while the algorithm was essentially unchanged from the previous year. It is worth noting that the top algorithm (U-net Multi-task Cascade (UMC)) requires noise-free images for training^34^. This approach is therefore not applicable on real cellular datasets, although there has been significant progress on using deep learning to denoise tomograms^29;30;36;37^.

In the following sections, we demonstrate on real datasets that DeepFinder successfully identifies macromolecules of varying shape and molecular weight in cellular cryo-ET datasets. In addition, we show that handling several molecular species simultaneously (i.e., multi-class strategy) is especially beneficial for identifying small macromolecules. The calculation of *F*_1_-scores usually relies on the availability of ground truth positions, as it is the case on synthetic datasets (SHREC’19 and SHREC’20), which are generally not available with experimental data. Instead of groundtruth, the *F*_1_ scores reported below are calculated with respect to the annotations provided by the experts which use a combination of template matching, visual curation, subtomogram averaging, and classification. A *F*_1_-score of 1 indicates that the performance of DeepFinder is the same as the approach supervised by the experts.

### DeepFinder robustly finds macromolecules of varying size in cellular tomograms

We quantitatively evaluated the performance of DeepFinder on several cellular tomograms. First, we explored the ability of our method to localize two ribosomal states (‘membrane-bound ribosomes’ and ‘cytosolic ribosomes’, denoted *mb-ribo* and *ct-ribo*, respectively), and we compared the results to those obtained with template matching (see Appendix A). In this study, we used a cryo-ET dataset (Dataset #2, see Methods) composed of 57 tomograms of *Chlamydomonas reinhardtii* cells and annotations of membrane-bound 80S ribosomes (~3.2 MDa) performed by experts^39^. In our experiments, we considered four classes to better detect the target macromolecules: membrane-bound ribosome (*mb-ribo*), cytosolic ribosome (*ct-ribo*), membrane, and background. The expert annotations contained positions of 8,795 *mb-ribos*. The training datasets corresponding to the *ct-ribo* and membrane classes were obtained by using semi-automated computational tools, without careful expert supervision (see Methods). The classes were highly imbalanced since the background class represents 99.5% of training voxels within the tomogram. As this scenario is common in cryo-ET, the training procedure (especially the generation of batches) needs to be adapted by integrating techniques such as re-sampling (see Methods). Figure 1B-C illustrates an example of our four-class segmentation and provides insight into the spatial distribution of two ribosomal states. When computing the overlap with the expert annotations, DeepFinder achieved higher scores than template matching. For *mb-ribos*, we obtained an *F*_1_-score of 0.86 for DeepFinder and 0.50 for template matching (Dataset #2, see Methods). DeepFinder detected *mb-ribos* that were not annotated by the experts. Several of these particles were false positives, but after a detailed inspection, we confirmed that many others are ribosomes that were missed or discarded during the annotation process.

To evaluate the performance of DeepFinder on smaller macromolecules, including both soluble and membrane proteins, we conducted experiments on two challenging datasets depicting pyrenoids (Dataset #3) (Fig. 3A-B) and thylakoid membranes (Dataset #4) (Fig. 3C-F) within *Chlamydomonas reinhardtii* cells^38;40^. In Dataset #3, DeepFinder was able to identify Rubisco holoenzymes with remarkable performance (Fig. 3A). The achieved *F*_1_-score (0.83) is similar to what was obtained with *mb-ribos* (0.86) (Dataset #2), even though the molecular mass of Rubisco (~540 kDa) is much lower than that of the 80S ribosome (~3.2 MDa). This strong performance was made possible by the large quantity of expert annotations (129,000 training particles from four tomograms). This result suggests that DeepFinder has the capacity to identify small macromolecular species with similar performance to larger complexes, provided that the training set size is large enough.

**Figure 3:**
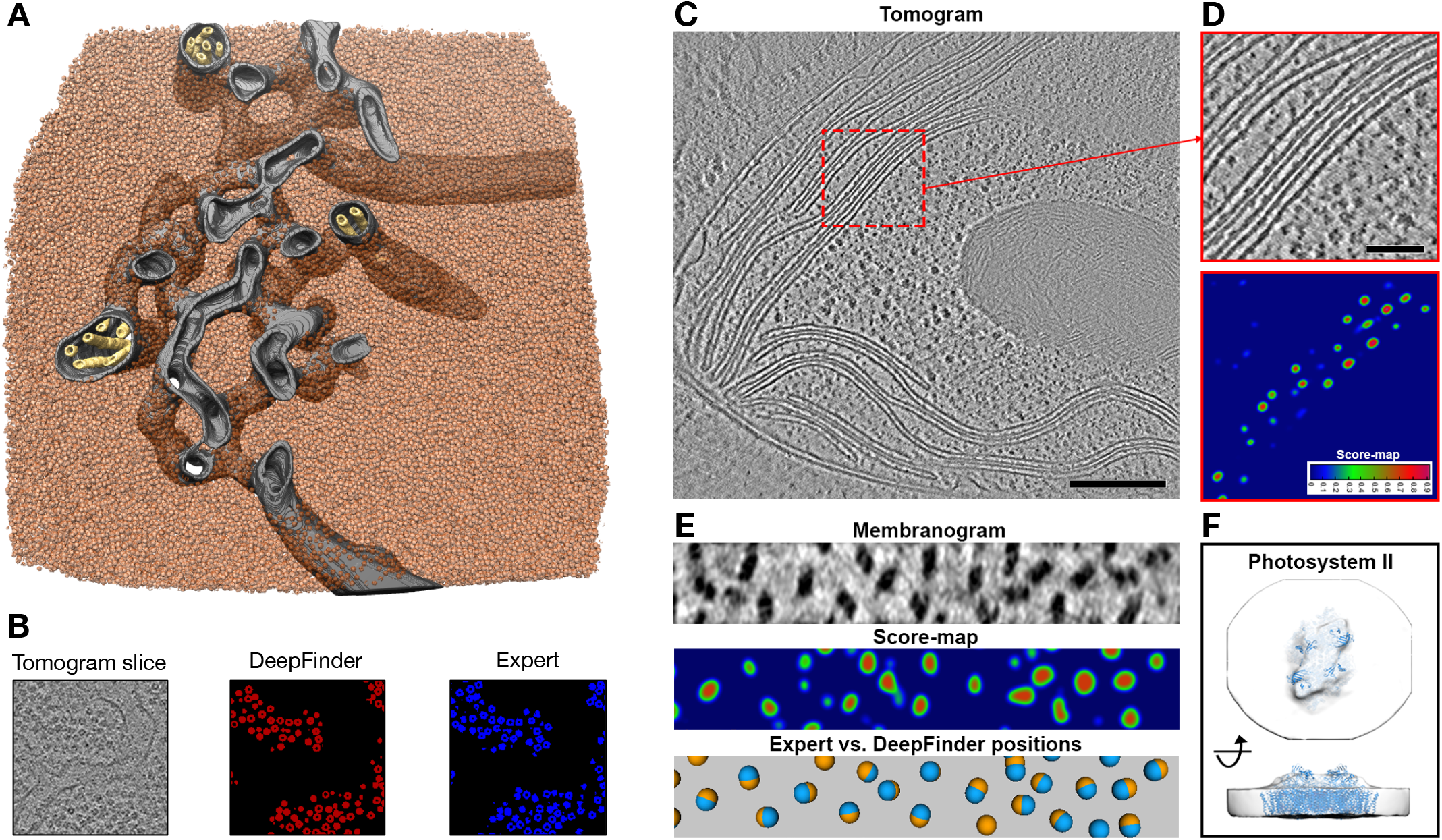
DeepFinder localizes small macromolecules in cellular tomograms. (A) Segmentation of a tomogram of the native Chlamydomonas pyrenoid (Dataset #3). Rubisco holoenzymes segmented by DeepFinder are displayed in transparent red. For visualization purposes, hand-segmented pyrenoid tubule membranes (gray and yellow) have been added. (B) Tomogram slice with corresponding DeepFinder segmentation and expert annotations. (C) Slice through a tomogram of the chloroplast within an intact Chlamydomonas cell (Dataset #4). (D) Zoomed-in view of the tomogram slice, with the associated score-map produced by DeepFinder. (E) The membranogram is a visualization approach^38^ where densities from the tomogram are projected onto the surface of a segmented thylakoid membrane, resulting in a topological view of the membrane surface. Below are the associated DeepFinder score-map and a visualization comparing expert manual annotations and DeepFinder localizations. (F) Subtomogram average (white) of photosystem II complexes, calculated from 246 particles identified by DeepFinder on a single test tomogram, fit with a PSII molecular structure (blue).

In our experiments on Dataset #4, DeepFinder was able to detect Photosystem II (PSII) dimers embedded within the thylakoid membranes (Fig. 3C-F). We trained DeepFinder on three tomograms using four particle classes that were manually annotated^38^: PSII dimers (~550 kDa), cytochrome b6f dimers (~220 kDa), unknown densities, and background. In Supplementary Table 1, we report the DeepFinder *F*_1_-scores when using the mono-class and multiclass strategies to detect PSII in the unseen test tomogram; the multi-class approach produced better results than the mono-class approach. PSII complexes were detected quite accurately in some membranes (as shown in Fig. 3E), which was confirmed by the high *F*_1_-score (0.737 for membrane 4 in Supplementary Table 1), whereas in other membranes they were not. Note that the number of particles per membrane used to compute scores is not the same in Supplementary Table 1). Overall, the mono-class approach with only PSII performed worse, in particular with respect to Recall, than the multi-class setting (0.505 versus 0.638 in Supplementary Table 1). This result is promising but limited due to the small training dataset size (298 PSII particles from 18 membranes) as well as the variable quality of the experimental data. Membranes where PSII complexes were poorly resolved had the most missed picks and lowest *F*_1_-scores. Nevertheless, the results are encouraging, and we are convinced that adding more particles to the training sets (as exemplified in the Dataset #3 experiment) would improve the capability of DeepFinder to reliably detect PSII and other membrane proteins.

Collectively, these experiments on soluble and membrane complexes of different sizes consistently indicate that the multi-class approach is a flexible and robust method to identify one target macromolecule. Moreover, spurious components such as gold beads or ice chunks can be typically annotated and included in the negative class (i.e., background) during the training task in order to reduce false positives, as illustrated in Supplementary Fig. 9.

### DeepFinder converts the 3D positions of annotated macromolecules into image targets for training

In the training stage (Stage I), DeepFinder converts the input 3D coordinates of macromolecules supplied by the experts into voxel-wise annotations (see Methods), avoiding the cumbersome and time-consuming manual annotation of voxels. It is worth noting that the DeepFinder software includes a graphical interface to help users explore the training tomogram and select the coordinates of macromolecules used to constitute the training dataset (see Supplementary Fig. 6). As converting the positions of macromolecules into voxel-wise annotations is not a trivial task, we propose two approaches with different levels of computing complexity. The first approach exploits the segmentation maps built from well-delineated shapes of macromolecules estimated by a subtomogram averaging procedure^41^. In the second approach, the shapes of macromolecules are approximated by 3D spheres (see Fig. 4). The sphere-based representation is appealing since neither information about the shape nor the orientation of the macromolecules are needed, and time-consuming subtomogram averaging steps are avoided. However, using 3D spheres instead of 3D shapes induces more “label noise”, which hinders optimization of the network. In our experiments on the SHREC’19 dataset, the sphere-based representation was able to provide good scores for large and medium sized macromolecules, as shown in Fig. 2C. For small and tiny macromolecules, the shape-based representation provided better results. For larger macromolecules, the “label noise” induced by the sphere-based representation is compensated by the high quantity of labeled voxels in the segmentation map (see Rolnick *et al*.^42^ for discussion). Small macromolecules have very few associated labeled voxels, and thus “label noise” more seriously hinders the learning procedure. In conclusion, the computationally cheap sphere-based representation is recommended, provided that the target macromolecule is not too small and topologically equivalent to a sphere. We draw your attention to macromolecule 4d8q, which is topologically similar to a 3D torus; in this particular case, the shape-based representation (*F*_1_ = 0.97) drastically outperformed the “compact” sphere-based representation *(*F*_1_ =* 0.13).

**Figure 4:**
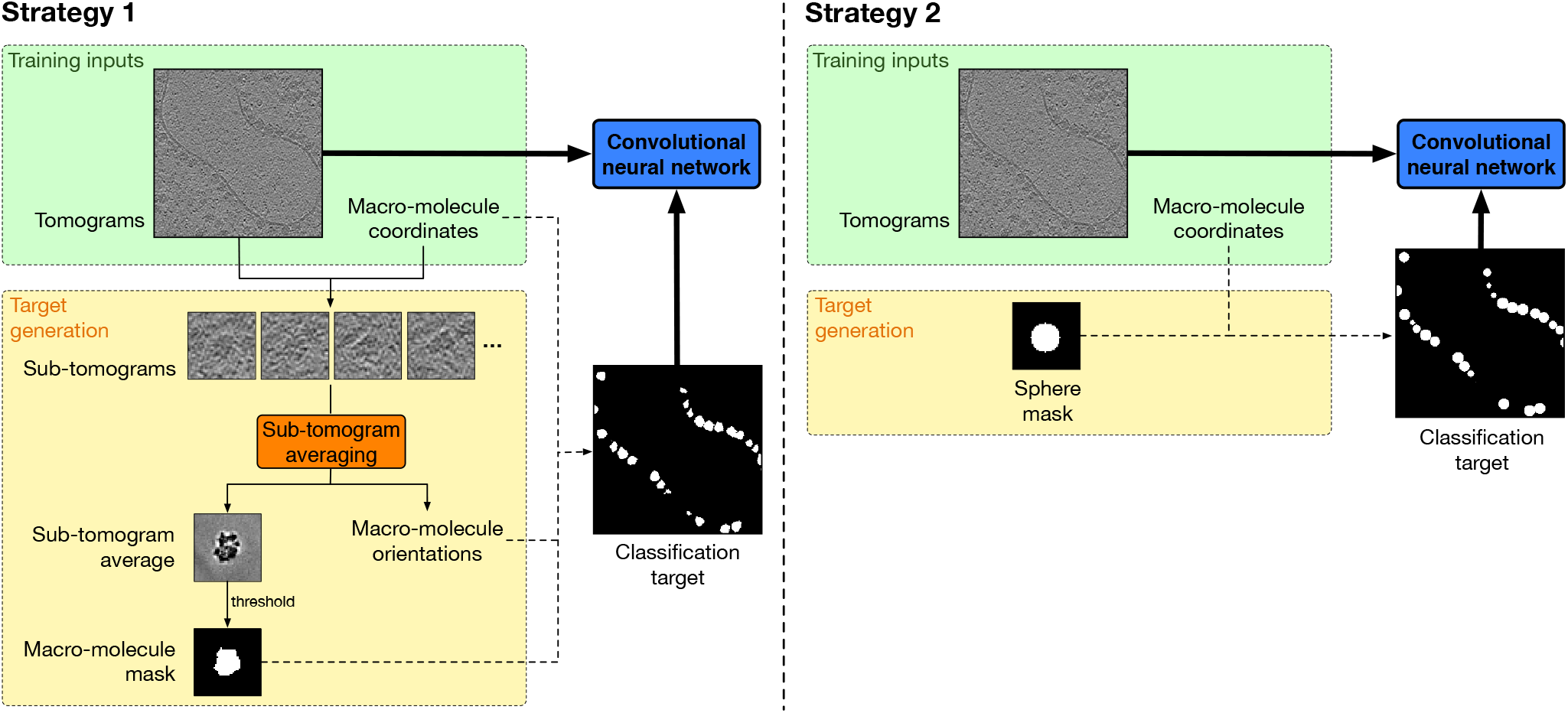
Target generation strategies for training. The DeepFinder software includes a graphical user interface (see Supplementary Fig. 6) to guide users throughout the learning procedure, from selecting the 3D coordinates of macromolecules to voxel-wise annotation. **Strategy 1 (left):** voxel-wise annotations are obtained from position-wise annotations using a subtomogram averaging procedure. Subtomogram averaging is a registration algorithm that produces higher resolution structures by averaging thousands of aligned subvolumes containing the molecular species. Subtomogram volumes are extracted at the annotated positions, aligned and finally averaged. The alignment procedure outputs the object orientations, while the averaging process provides a density map of the macromolecule with greatly reduced noise and missing wedge artifacts. From this density map, it is possible to create a binary mask of the macromolecule by thresholding the averaged subtomogram. The resulting mask is pasted into an empty volume at each annotated position with the estimated 3D orientation, and the resulting volume with well-delineated macromolecules is then used as a target to train the 3D CNN parameters. **Strategy 2 (right):** the macromolecule masks are approximated and replaced by sphere masks. Hence, the subtomogram averaging procedure is bypassed and the target generation process is much faster. However, the training targets contain more *”label noise”*.

### More time-consuming data annotation is required to localize small macromolecules

We analyzed the influence of the size of the training set on the performance of our method (see Supplementary Figs. 10-11). It is desirable to achieve good performance even if the training set size is small, as manual annotation of data is time consuming and requires considerable effort in 3D imaging. As shown in Supplementary Fig. 11 (synthetic SHREC’19 dataset), the performance of DeepFinder correlated with macromolecule size. For large and medium macromolecules, the performance is remarkably stable, even when using only one training tomogram (i.e., 206 macromolecules per class). For small and tiny macromolecules, the drop in performance is more significant, and larger training sets are required to get higher score values. This is not surprising since the number of labeled voxels associated with large macromolecules is high in the segmentation maps. Accordingly, less annotated macromolecules are needed. In Supplementary Fig. 10, note that for large and medium particles, we do not observe a monotonous increasing of the *F*_1_-score with respect to the number of training tomograms, mainly because of the stochastic nature of neural network training and the heterogeneous numbers of macromolecules in the training tomograms. On cellular cryo-electron tomograms, the *F*_1_-score is not significantly worse if we use a *mb-ribo* training dataset consisting of one-fifth of the complete annotated dataset (8,795 *mb-ribos*) (see Fig. 11). Together, these results suggest that DeepFinder does not require a large quantity of annotations for localizing large macromolecules, but more annotations are necessary to localize small macromolecular species. This is exemplified by the mixed results for ~550 kDa PSII with limited training data, but nice results for ~540 kDa Rubisco with extensive annotations.

### DeepFinder imitates expert annotations on experimental cryo-ET data

To carefully examine the performance of DeepFinder on cellular cryo-tomograms, we focused on ribosomes. These large and abundant macromolecules are relatively well localized with template matching, yielding abundant annotations for training and allowing a direct comparison of DeepFinder with the template matching results.

#### Analysis of score-maps

In Fig. 5, it is clear that the score-map of template matching (Fig. 5D) is much noisier than the score-maps generated by DeepFinder (Fig. 5B-C). Template matching produced high cross correlation scores at ribosome locations but also at false-positive locations containing other high contrast structures (e.g., cell membrane in Fig. 5D). Consequently, template matching is mainly used to crudely discard the voxels with no significant structural information. In the next step, the experts must apply post-processing techniques to separate the desired macromolecules from the false positives. Unlike template matching, DeepFinder produces less noisy score-maps, with meaningful score values in well-localized blobs.

**Figure 5:**
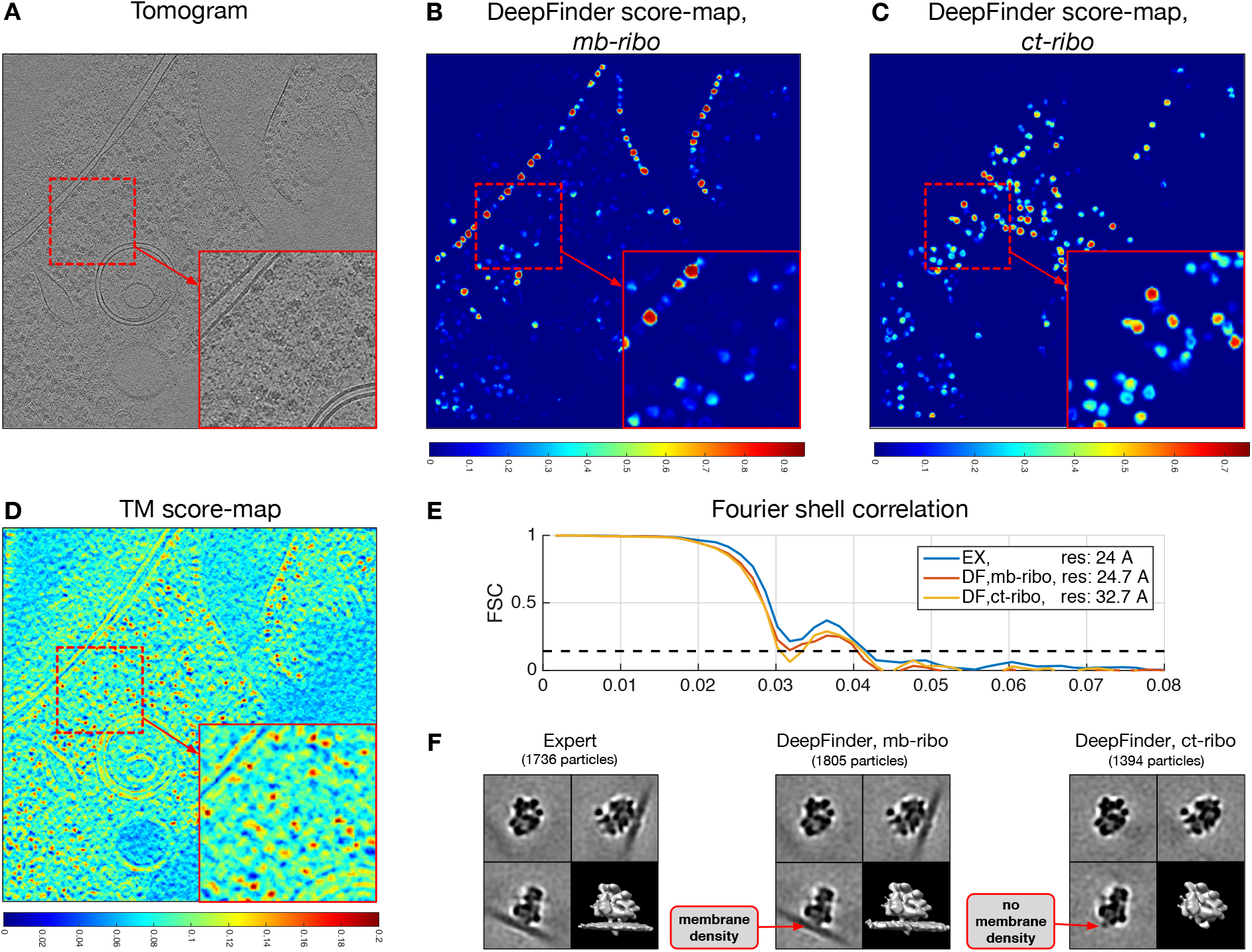
Comparison of score-maps obtained with template matching and DeepFinder and analysis of structural resolution through subtomogram averaging (Dataset #2). (A) Experimental cryo-electron tomogram depicting a *Chlamydomonas reinhardtii* cell (Dataset #2, see Methods). (B,C) Localizations of *mb-ribos* and *ct-ribo* particles with DeepFinder, and histograms of score-map local maxima. (D) Localizations of ribosomes with template matching (TM), and histogram of score-map local maxima. (E) Gold-standard FSC curves in each case with estimated resolutions. The resolution (24Å) corresponding to expert annotations (dedicated to *mb-ribo* particles) is very similar to the resolution (24.7Å) obtained with *mb-ribo* particles localized with DeepFinder. (F) Subtomograms obtained from expert annotations (left) and from particles localized with DeepFinder (*mb-ribos* (middle) and *ct-ribos* (right)). The average densities have been obtained with the same alignment procedure and parameter settings. For comparative visualization, the averages have all been low-pass filtered to 40Å resolution.

Furthermore, DeepFinder allows accurate localization of two the two ribosomal states in cellular cryo-ET data, as illustrated in Supplementary Fig. 6A. The *mb-ribos* (in blue) are primarily located close to cell membranes, whereas the *ct-ribos* (in yellow) are mainly located in the cytosol. This is confirmed by analyzing the Euclidean distances of *mb-ribo* and *ct-ribo* particles to the nearest cell membranes (see Supplementary Fig. 12). In the *mb-ribos*, we observed a significant mode located at 13.7 nm, which approximately corresponds to the radius of the ribosome; this suggests that the majority of the voxels classified as *mb-ribos* are very close or linked to the membrane. As for the *ct-ribo* class, the distribution was flatter, which is expected because the identified particles have no membrane density.

#### Analysis of consensus responses

For the next step of our evaluation, we compared the template matching and DeepFinder procedures by using the expert annotations as ground truth. In Supplementary Fig. 13, we plotted the *Recall, Precision* and *F*_1_-scores with respect to DeepFinder and template matching algorithm parameters. We focused on the thresholds used to detect objects with template matching (thresholding of score values) and DeepFinder (thresholding of object sizes). We obtained a maximum *F*_1_-score of 0.86 for DeepFinder and a *F*_1_-score of 0.50 for template matching (with no post-classification step, see Supplementary Fig. 7A) (Dataset #2). Furthermore, we also examined the complementarity between the two sets of *mb-ribo* macromolecules found by the experts and DeepFinder. In the following analysis, we denote the sets obtained by experts and DeepFinder as *S_E_* and *S_DF_*, respectively. While the overlap *S_E_* ⋂ *S_DF_* between both sets is substantial (1,516 particles), there is also a significant number of particles belonging to *S_E_* \ *S_DF_* (220 particles), i.e., the particles annotated by the expert but not found by DeepFinder, and to *S_DF_* \ *S_E_* (356 particles), i.e., particles found by DeepFinder but missed by the expert. We can benefit from the two complementary sets of particle positions to improve the overall validation rates. The union *S_E_* ∪ *S_DF_* of the two sets increases the list of potential *mb-ribo* macromolecules, for which a confidence level can be assigned to each set member depending on whether it belongs to *S_E_* ⋂ *S_DF_*, *S_DF_* \ *S_E_* or *S_DF_* \ *S_E_*. The particles belonging to *S_E_* ⋂ *S_DF_*, i.e., found by both methods, are very likely to be true positives. Meanwhile the particles belonging to *S_E_* \ *S_DF_* and *S_DF_* \ *S_E_* can be labeled as “suspicious” and require more investigation. These two non-union sets are relatively small, enabling assignment of the bulk high-confidence particles so the expert can focus on validating the remaining low-confidence particles. In this manner, it is possible to uncover inaccuracies in the expert annotations and refine the true-positive particle class, which can further improve the training performance of DeepFinder.

#### Analysis of structural resolution

DeepFinder achieves 3D structural resolution comparable to the expert annotations, as determined through subtomogram averaging. We analyzed the particles by computing subtomogram averages for each ribosomal state (*ct-ribos* and *mb-ribos*) (Fig. 5F). The input sub-volumes which served to compute subtomogram averages are extracted from tomograms taken in the testing set. We applied the same registration procedure (rotational matching^43^) before computing the averages. The resulting density maps of *mb-ribos* are composed of two parts: the ribosome density and the membrane density. As expected, the membrane density computed from the set *S_DF_* of *mb-ribo* particles is well defined, while there is no membrane density in the average computed from the set of *ct-ribo* particles. We noticed that the ribosome density is similar in the two ribosomal states, with a difference at the interface with the membrane.

Furthermore, we calculated the Fourier shell correlation (FSC) from the subtomogram averages for each ribosomal state (Fig.5E). For *mb-ribos*, the resolution was comparable between the expert annotations and DeepFinder (24Å and 24.7Å, respectively). The resolution was lower for the *ct-ribo* average (32.7Å) mainly because the *ct-ribo* annotations used for training, were not accurately curated by the experts. One important consideration when evaluating resolution is that the averages were generated from a limited number of particles (1,805 *mb-ribos* and 1,394 *ct-ribos)*. It worth noting here that the DeepFinder *ct-ribo* average contains 23% less particles than the DeepFinder *mb-ribo* average (1,394 vs. 1,805 particles) and is 8Å lower in resolution. Nevertheless, the goal of this exercise was not to produce the highest resolution average (which could require >10,000 particles), but rather to test how well DeepFinder can distinguish subpopulations, even when particle number is limited. We therefore carefully analyzed the difference between the two *mb-ribo* sets that were used to compute the subtomogram averages. We investigated the set *S_DF_* of *mb-ribo* particles found by DeepFinder and considered the two following hypotheses:

*H*_1_: The set *S_DF_* contains too many false positives.
*H*_2_: The set contains particles with a low quality (due to noise and blur), suggesting that our method found supplementary true *mb-ribo* particles that were missed or discarded during the supervised annotation.

To test these hypotheses, we aligned and averaged *mb-ribo* particles in the set *S_DF_* \ *S_E_*, i.e., found by DeepFinder but not present in the set of expert annotations. We observed that the corresponding density map depicts a ribosome structure (see Supplementary Fig. 14). Thus, hypothesis *H*_1_ is not likely. As shown in Supplementary Fig. 15, DeepFinder finds additional membrane-bound ribosomes, missed or discarded by experts (hypothesis *H*_2_). Therefore, it appears that the number of actual *mb-ribos* is higher than expected: in our test set, we detected +20.5% *mb-ribos* when compared to the *S_E_* set. This result confirms the benefit of combining several analysis methods in cryo-EM^44^. The collaboration between the expert processing chain and DeepFinder decreases the overall false negative rate and generates a larger set *S_E_* ∪ *S_DF_* of membrane-bound ribosomes. The extended set of particles is probably less homogeneous, as it contains supplementary particles which are noisier. Adding this set of additional particles in the average may blur the result and therefore degrade the structural resolution. Nevertheless, these particles not found by template matching should not be discarded *a priori*, as they provide a more complete picture of the cellular environment. In that sense^44^, the proposed collaborative strategy between DeepFinder and template matching may be relevant to better investigate cryo-ET data, in particular when attempting to accurately quantify the concentration or organization of macromolecules inside the cell.

## DISCUSSION

The analysis of cellular cryo-electron tomograms is very challenging because of the molecular crowding and high diversity of molecular species. Hence, we developed DeepFinder, consisting of training (Stage I) and analysis (Stage II) stages for supervised deep learning-based localization of multiple macromolecular species. In Stage II, the algorithm first produces a segmentation map where a class label is assigned to each voxel. The classes can represent different molecular species, states of a molecular species (e.g., binding states, functional states) or cellular structures (e.g., membranes). In the second step of Stage II, the segmentation map is used to extract the positions of macromolecules. To perform image segmentation, we use a 3D CNN with architecture and learning procedures that have been adapted for large datasets with unbalanced classes. With accessibility in mind, we developed graphically-assisted training strategies to guide the user in producing multiple segmentation maps from the spatial coordinates of macromolecules. In the first step of the analysis stage, DeepFinder classifies the voxels into different categories of macromolecules, including an optional ‘outlier’ class and a background class. In the second step of the analysis stage, a “mean-shift” algorithm^45^ is applied to cluster the voxels with the same label class. The detected clusters correspond to individual macromolecules, and the particle positions are defined by each cluster’s center of gravity. The “mean-shift algorithm has the advantage that it is controlled by only one parameter (depending on the macromolecule’s diameter) and that the number of clusters does not need to be known in advance (as opposed to the k-means algorithm, for instance).

To cope with the missing wedge effect (i.e., delocalization along Z-direction), the usual template matching procedure uses an isotropic template, along with missing wedge-constrained cross-correlation as a criterion. In contrast, DeepFinder is trained to predict a segmentation mask depicting the shapes of the particles, which is unaffected by the missing wedge (see Fig. 4). DeepFinder applies a series of linear (convolution) and non-linear operations (maxpooling, ReLU activation). All these cascaded operations are performed in real space and progressively modify the results in Fourier space, thereby reducing the influence of the the missing wedge. Even if some features are missing or distorted due to the missing wedge, particles can still be recognized based on the remaining intact features of the particles. Thereby, DeepFinder, like most CNNs, makes decisions based on features of various sizes in real space.

While identifying large macromolecular complexes (e.g., ribosomes) is a relatively feasible task, the big challenge in cryo-ET is to identify smaller complexes, with lower molecular weights. Unlike template matching and mono-class approaches, DeepFinder was able to find several macromolecules of varying size at once on competitive benchmarks (SHREC’19 and SHREC’20 challenges) and in real cellular datasets (e.g., 540 kDa Rubisco particles). On both synthetic and real datasets, the multi-class approaches consistently perform better than mono-class approaches, especially on small particles. A relatively good annotation of large molecular complexes improves identification of smaller macromolecules. Even though DeepFinder yielded the most promising results – among other competitive methods – for localizing small macromolecules, these results will be improved as larger quantities of training data are continually collected and annotated. While unsupervised learning holds great promise for the future, the current price to be paid for better results is to resort to supervised machine learning algorithms guided by time-consuming expert annotations.

Moreover, In the case of experimental data, annotations are a subset of the unknown ground truth. By investigating the difference between the two sets, we discovered that a subset of additional *mb-ribo* particles found by DeepFinder were missed or discarded during the supervised expert annotation. Meanwhile, DeepFinder skipped many of the false positive positions selected by template matching but filtered out later by the expert. Therefore, combining the two sets helps decrease the overall false negative rate while increasing the set of true positive particles (about 20% supplementary *mb-ribo* particles in our experiments). More generally, the combination of algorithms with different designs paves the way towards robust fully automated identification of macromolecules in cryo-ET^11^.

Currently, only a handful of different molecular species have been analyzed in cellular cryo-tomograms. These volumes contain rich information on the whole proteome, but many macromolecules are hardly discernable from noise and artifacts. To realize the potential of visual proteomics^49^ and study the interaction between several macromolecular species, more powerful pattern recognition techniques are required. We believe that deep learning is the right approach to address this challenge. Therefore, we have developed DeepFinder to efficiently identify molecular complexes with variable shapes and molecular weights. For the first time, to our knowledge, we showed quantitatively that we can obtain a structural resolution with a deep learning-based method that is similar to that obtained by expert curation. Moreover, once trained, DeepFinder is relatively fast compared to the common template matching and subtomogram classification pipeline. Larger datasets can be processed in one day with DeepFinder, while several macromolecular species are simultaneously identified. Furthermore, we provide an open-source implementation of DeepFinder, with a graphical user interface for both training and identifying macromolecules in cellular tomograms, aimed to be accessible for a broad base of users.

Future challenges include developing unsupervised deep learning methods^51^ and applying transfer learning^50^ in order to reduce human annotation effort and annotation errors. Transfer learning would allow download of pretrained networks that can directly applied to newly acquired data (see an example in Supplementary Fig. 16)). Other deep learning investigations that exploit the PDB (Protein Data Bank) and EMDB (Electron Microscopy Data Bank) databases are currently pursued to help expand the range of molecular species that can be identified in cellular tomograms.

## METHODS

### Description of eepFinder algorithm – Stage I (Training)

In the training stage (see Fig. 1A), DeepFinder parameters are learned from pairs of tomograms and their corresponding voxel-wise annotations. The underlying 3D CNN requires that each voxel is annotated as a member of a given macromolecular specie, or as background. While voxel-wise classification examples are naturally available for synthetic data, this is often not the case for experimental data. In our case, the experts only supplied the 3D coordinates of macromolecules of interest without labeling voxels, in several training tomograms. In practice, voxel-wise annotation cannot be performed manually in cryo-ET for two reasons: *i*) it is time consuming to individually label each voxel belonging to a 3D macromolecule; *ii*) noise and artifacts in the data make it difficult to distinguish macromolecule borders are barely visible.

To get voxel-wise annotations, we propose two strategies starting from 3D spatial coordinates of the particles:

- *Sphere-based annotation:* 3D spheres are positioned at each annotated location in the tomogram. The sphere radius should be similar to the size of the target macromolecule. As the experts generally know the size of the target macromolecule (e.g., 15 nm radius for ribosomes to 5 nm radius for small particles like Rubisco), this parameter can be readily input into DeepFinder. This strategy is easy to implement and is appealing in terms of computing time. The main limitation is that spheres may introduce significant “label noise” ^42^. The resulting errors in the annotations hinder the training procedure, and the segmentation network may have sub-optimal performance. Nonetheless, it has been shown that CNNs have a natural robustness to moderate levels of “label noise”^42^ which is also confirmed in our experiments with DeepFinder.
- *Shape-based annotation:* To reduce the amount of “label noise”, we propose a second strategy to convert macromolecule coordinates into voxel-wise annotations. Instead of using spheres, we generate the target by using the structural shape of the macromolecule. These targets are more difficult to obtain, because information on the shape and the orientation of the macromolecules are needed. In order to obtain this information, we exploit the commonly used subtomogram averaging procedure^41^ (see Fig. 4). While this strategy provides more accurate targets, it still induces “label noise”, albeit less than when using spheres. Indeed, as we project the same average shape (computed with subtomogram averaging)) into the tomogram several times at different positions into the tomogram for training, all structural details of individual macromolecules are not captured.

In our experiments, all results have been obtained by using shape-based annotations, except for the comparison between sphere-based and shape-based annotations (Fig. 2C). Computing times for Stage I and Stage II are reported in Supplementary Table 2, and are relatively small when compared to template matching. For instance, if we consider a set of 10 tomograms, the computing times are 300 hours (10 × 30 hours) hours hours with template matching, while it is 23.30 hours (20 hours (training) + 10 × 20 min (inference)). This estimation does not take into account the time needed to annotate the data (for DeepFinder) nor the time to generate the template and classify the results (for template matching). Moreover, as the DeepFinder training expands and becomes more generalized, less training is required to analyze new data, further increasing its speed advantage over template matching. Note that the shape-based target generation takes more computing time than the sphere-based strategy, because it relies on a sub-tomogram alignment routine (with no classification procedure) performed on sub-resolved (binned) tomograms of size 47 × 47 × 47 voxels.

#### Training datasets and batch generation

Due to memory limitations, it may not be feasible to load the whole tomogram set with the corresponding targets during training. Instead, for each training iteration, we sample a group of smaller 3D patches to constitute a “batch”. The patch size should be large enough to capture sufficient context information. For a macromolecule with a 10 voxel radius, we choose a patch size of 56 × 56 × 56 × voxels. Our implementation is such that only the current batch is loaded into memory. This helps to handle large datasets with limited computational resources, allowing DeepFinder to be applied on lower-end consumer GPUs with less RAM.

Another problem is the high imbalance between class labels. Because of the small size of the macromolecules compared to the tomogram size, more than 99% of voxels are annotated as “background”. This causes the trained network to be skewed towards the over-represented class. Therefore, we “guide” the sampling procedure by selecting patches only at annotated positions such that each patch contains at least one macromolecule. An additional benefit of sampling only at annotated positions is that the amount of false negatives is reduced (here the negative class is the background). It is indeed common that annotations are not exhaustive. Therefore false “background” labels remain at missed macromolecule positions, contributing to increased “label noise”. The proposed patch sampling procedure does not discard all false negatives (e.g., a false positive could be neighboring a true positive), but the number should be relatively small at the end.

An additional problem is the imbalance between competing macromolecule classes: for instance, 20 macromolecules of class #1 (e.g., *mb-ribo* particles) versus 100 macromolecules of class #2 (e.g., *ct-ribo* particles). To reduce this imbalanced class effect, we apply a bootstrapping algorithm (i.e., resampling), so that the distribution of the positive classes in a batch will be uniform. This stochastic resampling procedure is effective for sampling the under-represented classes more frequently than the over-represented classes.

It is also common in deep learning to use “data augmentation” to artificially increase the size of training sets. The idea is to apply geometric transform to training images. In our approach, we implement “data augmentation” by applying a 180° rotation with respect to the microscope tilt-axis, to each training example. Nevertheless, we do not use “mirror” operations or geometric deformations because the macromolecule structure (including its chirality) is the main clue for detection. Also, we do not use random rotations because of the well-determined orientation of missing wedge artifacts, which is preserved when applying 180° rotations with respect to tilt-axis. Finally, we apply random shifts to improve invariance to translations.

#### Optimization

In our experiments, DeepFinder has been computationally trained with the ADAM algorithm, chosen for its good convergence rate^?^, by setting the learning rate to 0.0001, the exponential decay rate to 0.9 for the first moment estimate, and to 0.999 for the second moment estimate. The batch size was set to 25 and the patch size to 56 × 56 × 56 voxels. No regularization (e.g., *L*_2_ regularizer, “drop out”) was needed for processing the datasets. We implemented both the categorical cross-entropy (as implemented in Keras) and the Dice^?^ loss functions. On the synthetic SHREC’19 dataset, Dice loss function was able to better localize the smallest macromolecules. However, both loss functions yielded similar results on real cryo-ET data (localization of *mb-ribo* and *ct-ribo* particles).

### Description of DeepFinder algorithm – Stage II (Analysis)

We describe below the two steps of the analysis stage of DeepFinder algorithm (see Fig. 1A).

#### Analysis stage (Stage II): Step #1 – Multi-class voxel-wise classification

The objective is to provide a segmentation map for which each voxel is assigned to a class label (representing a macromolecular species). The architecture of DeepFinder is based on U-net, an “encoder-decoder” type network designed for segmenting images in an end-to-end manner. U-net is an extension of the fully convolutional networ^k16;20^, which achieves multi-resolution feature representation and produces high-resolution label maps. The DeepFinder architecture consists of a downsampling path (i.e., encoder) needed to generate global information and an up-sampling path (i.e., decoder) used to generate high-resolution outputs? ?, i.e., local information (see Supplementary Fig. 8). Down-sampling is performed with max-pooling layers (factor 2) and up-sampling with up-convolutions^16^ (sometimes called “backward convolution”), which is essentially a trained and non-linear up-sampling operation. Combining global and local information is performed by concatenating features at different spatial resolutions. The features are then processed with the convolutional layers of the up-sampling path. The fully convolutional nature of DeepFinder allows one to use various input patch shapes, with the constraint that the patch size must be a multiple of four, because of the two down-sampling stages. Large tomograms can be processed by using an overlap-patch strategy. Unlike Milletari *et al*.^?^, DeepFinder is not that “deep”, since we found that using more than two down-sampling stages does not increase the classification results. Also, we used only 3 × 3 × 3 filter sizes as described by Simonyan *et al*.^?^. The rationale behind this choice is that two consecutive 3 × 3 × 3 filters mimic a larger 5 × 5 × 5 filter but with fewer parameters. Training is then faster and easier and requires less memory. An important concept in neural architectures is the receptive field of deepest neuron layers. This determines the size of the spatial context to be used to make decisions. Considering a large spatial context is essential to handle macromolecule classes involving interactions with the environment, for instance interactions with cellular membranes. It is established that adding convolutional layers after down-sampling operations is appropriate to enlarge the spatial context^?^ . Accordingly, we added two supplementary convolutional layers in the lowest stage of our architecture. In the end, the receptive field size of our network is 48 × 48 × 48 voxels. To complete the description, we use rectified linear units (ReLU)^15^ as activation functions for every layer except the last one which uses a *soft-max* function. While ReLU is a popular choice to tackle non-linearities in the network, the *soft-max* function is necessary in order to interpret the network outputs as pseudo-probabilities for each class. In summary, step#1 is capable of robustly classifying the cryo-ET tomogram into *N* classes with a high accuracy.

#### Analysis stage (Stage II): Step #2 – Clustering for macromolecule localization

Given the multi-class voxel-wise classification map and classification errors, the objective of step #2 is to estimate the spatial coordinates of each particle of a given macromolecular species. The voxel labels should be spatially clustered into distinct 3D connected components, each cluster corresponding to a unique particle. Because of noise, non-stationarities in the background, and artifacts in the tomogram, step #1 generates isolated labels or very small groups of voxels, as well as groups that contain different labels. Post-processing is then necessary to aggregate neighboring voxels into 3D connected components and to assign a unique label to each component (or cluster). The clusters significantly smaller than the size of target particles, are considered as false positives and are discarded. The centroids of meaningful clusters are computed to determine the location of the particles. As the centroid is computed by uniformly averaging the coordinates of cluster voxels, we are able to numerically produce positions with sub-voxel precision. As several voxel labels can be spatially grouped in a given cluster, the most frequent label is assigned to the detected particle. To address this task, we used the popular “mean-shift” clustering algorithm^45^ The main advantage of “mean-shift” is that it is controlled by only one parameter, commonly called the bandwidth, which is directly related to the average size of macromolecules. The K-means algorithm was not considered further since the number of clusters must be provided as input parameter by the user.

### Description of Datasets

#### Description of synthetic data (Dataset #1)

The synthetic dataset was generated by the cryo-Electron Microscopy group of Utrecht University for the challenge “SHREC 2019: Classification in Cryo-Electron Tomograms”, organized for the Eurographics Workshop on “3D Object Retrieval”^33^. The dataset has been especially designed to objectively evaluate the performance of localization and classification algorithms in cryo-electron tomography. The dataset was created as follows^33^ *i*) tomogram density maps were generated using Protein Data Bank (PDB) structures (identifiers: 1bxn, 1qvr, 1s3x, 1u6g, 2cg9, 3cf3, 3d2f, 3gl1, 3h84, 3qm1, 4b4t and 4d8q); *ii*) a series of projection images was generated with a ±57° tilt-range and a 3° tilt-step; *iii*) projection images were degraded by adding noise and contrast transfer function such that signal-to-noise ratio was 0.02; *iv*) the tomogram was reconstructed using weighted back-projection. The resulting tomograms have a size of 512 × 512 × 512 voxels and a voxel size of 1 nm.

In our experiments, we split the dataset #1 into training, validation and test sets. The training set was composed of 8 tomograms annotated with a total of 19,956 macromolecules. The validation set was composed of one tomogram annotated with a total of 2,490 macromolecules. The test set was composed of one tomogram annotated with a total of 2,540 macromolecules. The training was performed until convergence, which took on average 10,000 iterations and 33 hours of computation using the Dice loss function on a Nvidia Tesla K80 GPU.

#### Description of experimental cryo-ET data (Dataset #2)

The real cryo-ET dataset is composed of 57 tomograms of *Chlamydomonas reinhardtii* cells, and was used for membrane-bound 80S ribosome (*mb-ribo*) annotation. To produce the annotations, the experts used template matching, and then filtered out the false positives by applying the CPCA subtomogram classification algorithm^10^, and careful visual inspection (see illustration in Supplementary Fig. 7). To reduce the computational cost, the tomograms were binned to a voxel size of 13.68Å with tomogram dimensions of 928 × 928 × 464 voxels. Tilt range was ±60° with an increment of 2°.

As starting point, Dataset #2 was annotated by localizing 8,795 *mb-ribo* particles under expert supervision. Then, we introduced two additional classes (without expert supervision): the cytosolic ribosomes (*ct-ribos*) class and the *membrane* class. Annotations for these two additional classes were obtained by applying semi-automatized computational tools to tomograms. First, we selected members of the *membrane* class by employing an algorithm dedicated to cell membrane segmentation?. Meanwhile, we obtained training (*ct-ribo*) particles by applying template matching and selecting the most isolated candidates, located further than 273.6Å (i.e., the ribosome diameter) from the segmented membranes. The motivation behind adding new classes to the available annotations was twofold: on the one hand, we wanted to demonstrate the ability of DeepFinder to localize and identify multiple molecular species on real data with one pass; on the other hand, we noticed that the multi-class approach tends to improve the discriminating power of CNN, therefore better finding *mb-ribo* particles in experimental tomograms. When trained with only the *mb-ribo* examples, DeepFinder additionally finds unwanted cytosolic ribosomes. By considering the *ct-ribo* class in the training, we encourage the network to better discriminate both ribosome subclasses (corresponding to binding states). Note that these annotations of membrane components and cytosolic ribosomes have been obtained without the supervision of an expert. Therefore more errors are expected for these two classes when compared to the *mb-ribo* examples, reliably annotated by the experts.

Dataset #2 was arbitrarily split into training, validation and test sets. The training set was composed of 48 tomograms annotated with 6,834 *mb-ribos* and 6,687 *ct-ribo* particles. The validation set was one tomogram annotated with 222 *mb-ribos* and 254 *ct-ribo* particles. The test set was composed of eight tomograms annotated with 1,736 *mb-ribos* and 2,594 *ct-ribo* particles. In our experiments, we considered “background” and three classes: *mb-ribo*, *ct-ribo* and *”membrane”*. The “background” voxels are those that do not belong to any of the other three classes. Training was achieved using the categorical cross-entropy as a loss function (we got very similar results with the Dice loss function). The training was performed for 6,000 iterations and took 20 hours of computation on a Nvidia Tesla K80 GPU (see plots of loss evolution during training in Supplementary Fig. 17).

#### Description of experimental pyrenoid cryo-ET data (Dataset #3)

The dataset is composed of five tomograms of the Chlamydomonas pyrenoid, which have been annotated with a total of 176,229 Rubisco holoenzymes. The tomogram size is 928 × 928 × 464 voxels, the voxel size is 13.68Å, and the tilt range is ±60° with an increment of 2°. The expert annotations were obtained with template matching (excluding hits outside the pyrenoid matrix using a manually segmented mask), followed by subtomogram classification using CPCA clustering^10^. For more details about the dataset and the annotations, see^40^.

Dataset #3 was split into training and test sets. The training set was composed of four tomograms annotated with 129,662 Rubisco complexes. The test set was composed of one tomogram annotated with 46,567 Rubisco complexes. The training was performed for 20,000 iterations and took 35 hours on a Tesla K80 GPU. The loss curve converged after 10,000 iterations, suggesting that only half the time (i.e., 17.5 hours) would have been sufficient to obtain the presented results.

#### Description of experimental thylakoid cryo-ET data (Dataset #4)

The dataset is composed of four tomograms of Chlamydomonas thylakoid membranes, annotated with a total of 637 photosystem II (PSII) complexes. The bin4 tomogram size is 928 × 928 × 464 voxels with a voxel size of 13.68Å, and the tilt range is ±60° with an increment of 2^°^. The expert annotations were obtained by manual particle picking using the “membranogram” approach^38^. For more details about the dataset and the annotations, see^38^.

Dataset #4 was split into training, validation and test sets. The training set was composed of three tomograms annotated with 298 PSII complexes. The validation set was a region from one of the training tomograms annotated with 143 PSII complexes. The distance between training and validation particles was large enough to avoid overlap. The test set was composed of one tomogram annotated with 196 PSII complexes. The training was performed for 5,000 iterations and took 3.5 hours of computation on a Nvidia Quadro RTX 5000 GPU.

### Template Matching method

The template matching results on the synthetic SHREC’19 dataset were provided by courtesy of Utrecht University (Department of Information and Computing Sciences, Department of Chemistry) and were obtained as follows: *i*) template matching was applied for each class. As the particles have been simulated, we know the true number of candidates for each tomogram: 2,500 particles/12 classes = 211. Then, the top 211 candidates for each class were sequentially selected and extracted, from the largest macromolecules to the smallest ones, in a way that all these candidates do not overlap with the already extracted candidates.

We used the template matching algorithm implemented in the PyTom toolbox^35^ to analyze Dataset #2 (*Chlamydomonas reinhardtii* cells).

### Evaluation procedure

We used the *F*_1_-score to assess localization performance. The *F*_1_-score (also known as Dice coefficient) is the harmonic mean of *Precision* and *Recall* that depend on the number of true positives (TP):

- *Recall* (also known as sensitivity):

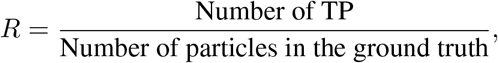
- *Precision* (also known as positive predictive value):

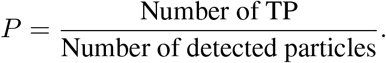

The *F*_1_-score, defined as follows

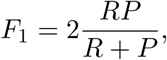

allows one to evaluate performance by considering a single value.

Moreover, for each test tomogram, bounding boxes were placed at each true location. These bounding boxes were of same size as the target macromolecule, and a detected particle was considered to be a TP if the estimated centroid was located within the bounding box.

Dataset #1 was very helpful to objectively study the performance of the algorithms. In the case of real cryo-ET data (Dataset #2), the spatial coordinates of *mb-ribo* particles supplied by the experts were used place bounding boxes and to compute *F*_1_-scores. Unlike Chen *et al*. ^31^, we also analyzed the score distributions and subtomogram averages. The resolution of obtained subtomogram averages was estimated with the commonly-used gold standard Fourier shell correlation (FSC) with a 0.143 threshold.

## Code availability and implementation details

Each step of DeepFinder shown in Fig. 1A can be executed with scripts using the API (examples are provided) or with a graphical user interface. These steps may also be embedded in other workflows, e.g., if the user needs only the segmentation step. To implement DeepFinder, we used Keras [http://keras.io], an open-source toolbox written in Python and using the TensorFlow framework. The code can be downloaded for free from the following GitLab website: [https://gitlab.inria.fr/serpico/deep-finder] and includes online documentation: https://deepfinder.readthedocs.io/en/latest/.

The execution times of Stage II to process a 928 × 928 × 464 tomogram are 6 min for step #1 (multi-class voxel-wise classification) and 15 min for step #2 (clustering), respectively.

All training procedures were achieved using a Nvidia Tesla K80 GPU, running Cuda 8 and cuDNN 6. Below, we display the memory consumption of DeepFinder for different training parameters.

| Patch size (voxels) | Batch size |  |
| --- | --- | --- |
|  | 15 | 25 |
| $40 \times 40 \times 40$ | 2.56 GigaBytes | 3.76 GigaBytes |
| $56 \times 56 \times 56$ | 6.16 GigaBytes | 9.86 GigaBytes |

We used the PyTom toolbox^35^ (template matching, subtomogram averaging, fast rotational matching^43^ for subtomogram alignment). The alignment has been performed with respect to a reference template. We used Chimera^?^ for 3D visualization purposes. TomoSegMemTV? was used to annotate the “membrane” class.

## Data availability

The synthetic dataset is available on the website of the SHREC’19 challenge (http://www2.projects.science.uu.nl/shrec/cryo-et/2019/). A tomogram from the experimental dataset of *Chlamydomonas reinhardtii* cells^39?^ can be found in the Electron Microscopy Data Bank (EMDB) (https://www.ebi.ac.uk/pdbe/entry/emdb/EMD-3967) under accession number EMD-3967. Additional tomograms depicting Thylakoids can be downloaded from EMDB under accession numbers EMD-10780, EMD-10781, EMD-10782, and EMD-10783.

## Acknowledgements

This work was jointly supported by the Fourmentin-Guilbert Foundation, and Région Bretagne (Brittany Council). Experiments on real data (courtesy of the MPI Biochemistry, Martinsried, Germany) were performed on the Inria Rennes computing grid facilities partly funded by France-BioImaging infrastructure (French National Research Agency - ANR-10-INBS-04-07, “Investments for the future”). L. Lamm, W. Wietrzynski, T. Peng, and B. Engel were supported by DFG grant EN 1194/1-1 as part of FOR 2092, The Munich School of Data Science, and Helmholtz Zentrum München.

We would like to thank F. Förster and M. Killinger for fruitful discussions about cryo-ET data analysis, and deep learning applied to large 3D volumes analysis, respectively.

We thank the organizers of the SHREC’19 challenge for helpful assistance and for providing the template matching results: I. Gubins and R.C. Veltkamp (Utrecht University, Department of Information and Computing Sciences), G. van der Schot and F.G. Forster (Utrecht University, Department of Chemistry).

Finally, we thank S. Prima for careful reading of the paper and valuable suggestions and comments.

## Author contributions

E. Moebel designed and implemented the presented DeepFinder method, carried out the biocomputing experiments, and wrote the manuscript. C. Kervrann supervised the project, was in charge of overall direction and planning, and co-wrote the manuscript. D. Larivière, E. Fourmentin, and C. Kervrann devised the project and the main conceptual ideas. A. Martinez-Sanchez also assisted the conceptualization. B. Engel and W. Baumeister facilitated access to datasets. B. Engel, S. Albert, W. Wietrzynski, and S. Pfeffer provided the Chlamydomonas reinhardtii datasets and annotations (Datasets #2, #3 and #4). A. Martinez-Sanchez, J. Ortiz and B. Engel conceived experiments on real datasets. L. Lamm, R. Righetto, W. Wietrzynski, and T. Peng performed experiments on datasets depicting thylakoid membranes and pyrenoid matrices within vitreously-frozen *Chlamydomonas reinhardtii* cells. B. Engel substantively revised the manuscript. All authors provided critical feedback and helped shape the research, analysis and manuscript.

## Note

Through the LifeExplorer project, the Fourmentin-Guilbert Foundation has been a pioneer in approaching the structural modeling and visualization of entire cells. It is expected from the creation of interactive 3D avatars of cellular environments, bridging from the level of atoms to the level of cells, that the rules governing the spatiotemporal organization of the cytoplasm could be revealed. Such an approach requires to make an inventory and a cartography of all the components constituting a single cell.

The technique of choice for such a mapping is cryo-electron microscopy applied on frozen but intact cells. For years, the Fourmentin-Guilbert Foundation has supported the Max Planck Institute of Biochemistry, headed by W. Baumeister whose team has been capable of delivering whole cryo-electron tomograms of E. coli cells at an unprecedented resolution.

The next big challenge, still an ongoing challenge, was to recognize macromolecular components within the tomograms. Most of the effort of the scientific community was put on the identification of the ribosomes thanks to a methodology relying on single-particle analysis and template matching and giving impressive results. However, it is likely that such an approach will be limited to large macromolecules like the ribosomes. Facing this challenge, the Fourmentin-Guilbert Foundation has solicited the research group headed by C. Kervrann to develop alternative recognition methods having the potential to help the *in-situ* identification of the thousands of proteins left in the dark. The Serpico Project-Team has then developed and compared with well-established methods new approaches based on deep learning and capable of identifying and counting the “gold standard” ribosomes within a tomogram. Theses methods, as an alternative to template matching, should also have the potential to apply on particles smaller and rounder than ribosomes.

## Supplementary information

**Table 1:** Comparison of *F*_1_-scores for the mono-class and multi-class strategies applied to detect PSII membrane complexes embedded within native thylakoid membranes.

| Membranes |  | M1 | M2 | M3 | M4 | M5 | M6 | M7 | M8 | M9 | Global |
| --- | --- | --- | --- | --- | --- | --- | --- | --- | --- | --- | --- |
| **Mono-class PSII** | $F_1$-score | 0.557 | 0.465 | 0.533 | 0.625 | 0.789 | 0.516 | 0.571 | 0.588 | 0.286 | **0.566** |
|  | Precision | 0.710 | 0.476 | 0.800 | 0.625 | 0.789 | 0.727 | 0.556 | 0.500 | 0.400 | **0.643** |
|  | Recall | 0.458 | 0.455 | 0.400 | 0.625 | 0.789 | 0.400 | 0.588 | 0.714 | 0.222 | **0.505** |
| **Multi-class PSII** | $F_1$-score | 0.632 | 0.582 | 0.714 | 0.737 | 0.696 | 0.250 | 0.562 | 0.182 | 0.571 | **0.619** |
|  | Precision | 0.638 | 0.485 | 0.625 | 0.636 | 0.593 | 0.750 | 0.600 | 0.250 | 0.800 | **0.601** |
|  | Recall | 0.625 | 0.727 | 0.833 | 0.875 | 0.842 | 0.150 | 0.529 | 0.143 | 0.444 | **0.638** |

**Table 2:** Computing time obtained for processing Dataset #2, composed of 57 tomograms of size 928 × 928 × 464 voxels. Our 32-core CPU cluster only needs 37 minutes for aligning and averaging 7,642 sub-resolved tomograms of size (47 × 47 × 47) voxels.

|  |  | Computing time | Hardware |
| --- | --- | --- | --- |
| **Stage I** | Target generation, sphere-based | 3sec / tomogram | 2.9GHz Intel Core i7 |
|  | Target generation, shape-based | 35min + (3sec/tomo) | 32-core CPU cluster |
|  | Training (until convergence) | 15 hours | Tesla K80 GPU |
| **Stage II** | Analysis |  |  |
|  | Step 1: segmentation | 20min / tomogram | Tesla K80 GPU |
|  | Step 2: clustering | 5min / tomogram | 2.9GHz Intel Core i7 |

**Figure 6:**
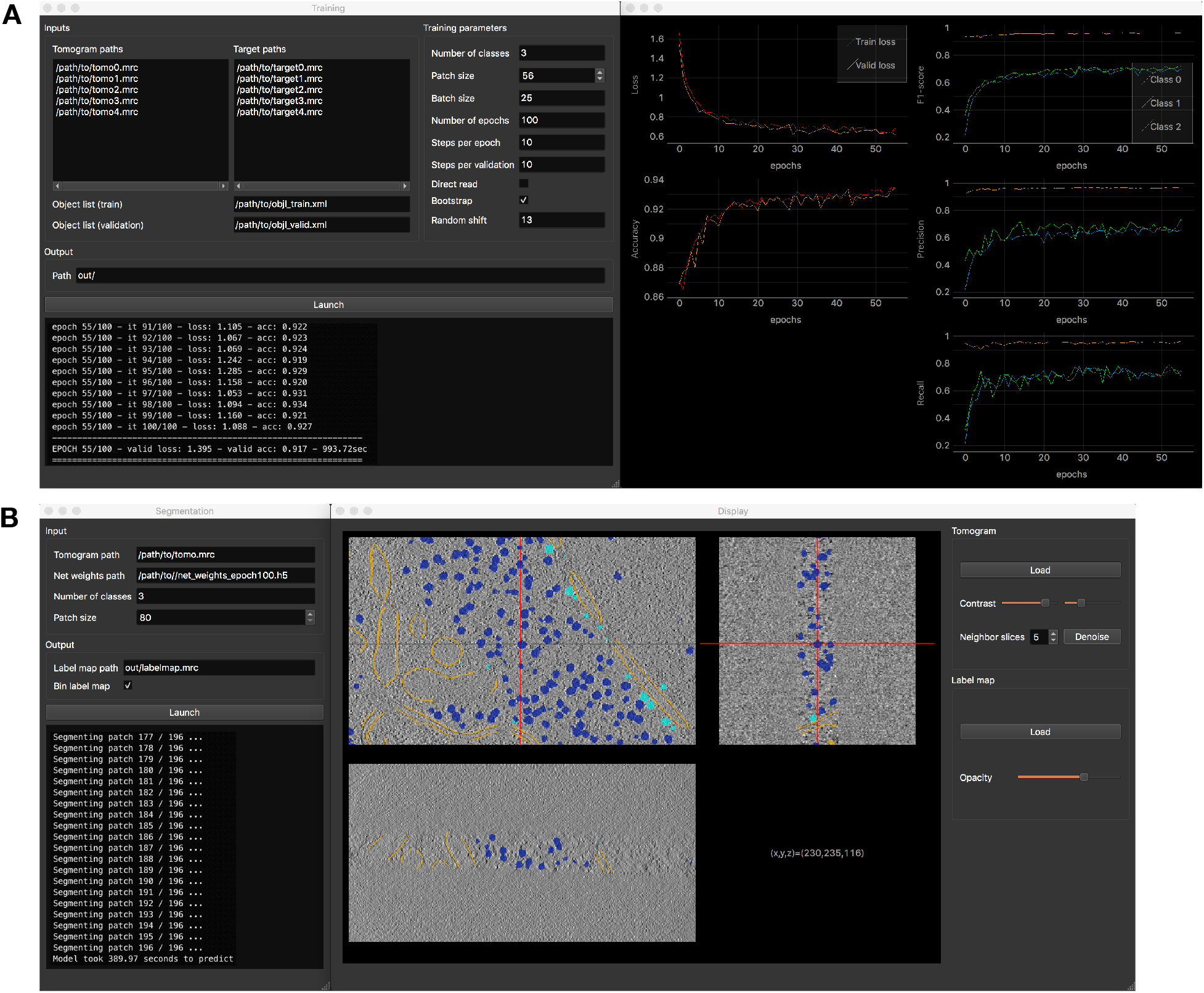
DeepFinder Graphical User Interface (GUI). (A) Training interface composed of a first window for parametrizing the procedure and a second window for displaying the training metrics in real-time. (B) Segmentation interface which also opens a data visualization tool. This tool allows one to explore the tomogram with superimposed segmentations. In addition, DeepFinder also incorporates interfaces for tomogram annotation, target generation and clustering (see the documentation at https://gitlab.inria.fr/serpico/deep-finder for more information).

**Figure 7:**
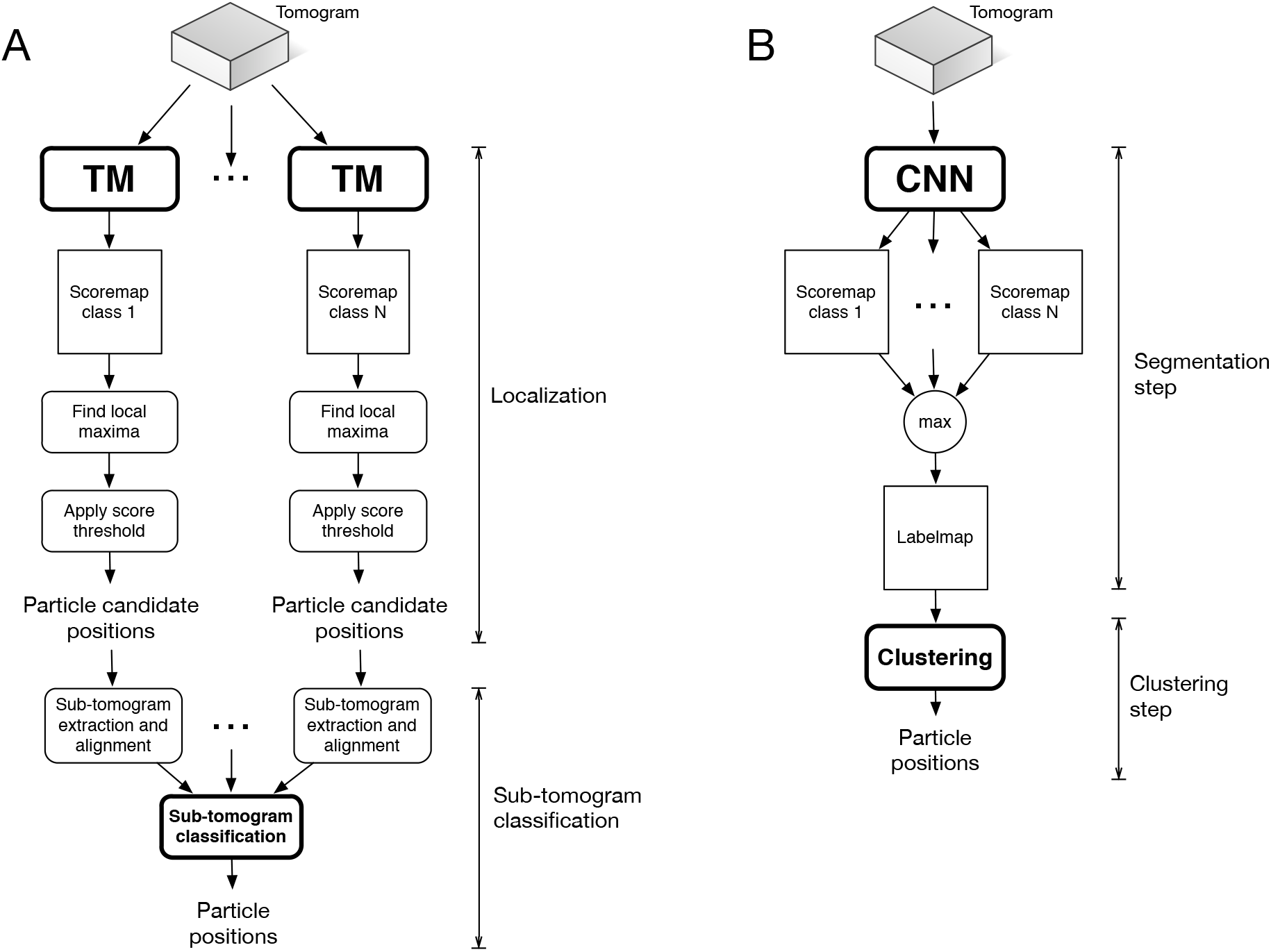
Two workflows for macromolecule localization in cryo-ET. (A) Conventional processing pipeline based on template matching. (B) DeepFinder (analysis stage): a multi-class approach able to localize several particles of different macromolecular species in one pass.

**Figure 8:**
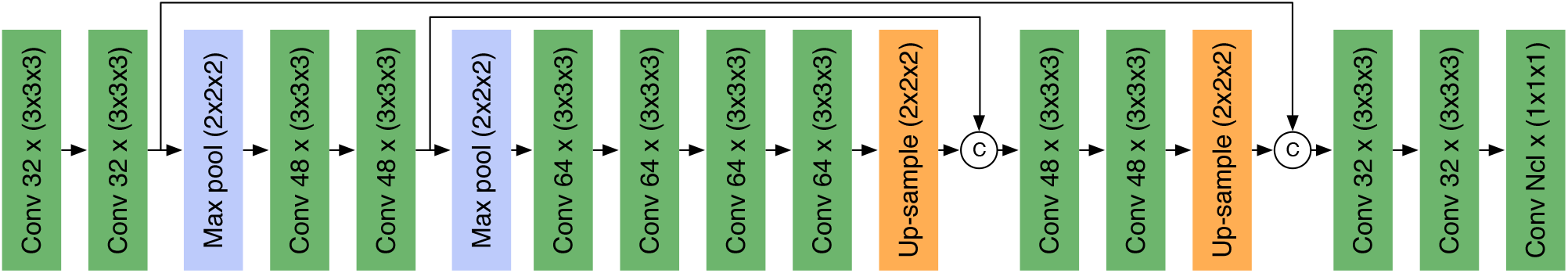
U-net architecture used in DeepFinder. The architecture adopts the encoder-decoder paradigm, which allows to produce an output volume having the same size as the input volume. Each green box represents a convolutional layer. The number of filters *n* and the filter size *s* is labeled as *n* × (*s* × *s* × *s*). All convolutional layers are followed by a ReLU activation function, except the last layer, which uses a soft-max function. The up-sampling is achieved with up-convolutions (also called “backward-convolution”). Combining feature maps from different scales is performed by concatenation along channel dimension. In the end, the total number of architecture parameters is approximately 903k. More precisely, this number depends slightly on *N_cl_*, the number of classes: 902928 + *N_cl_* × 33.

**Figure 9:**
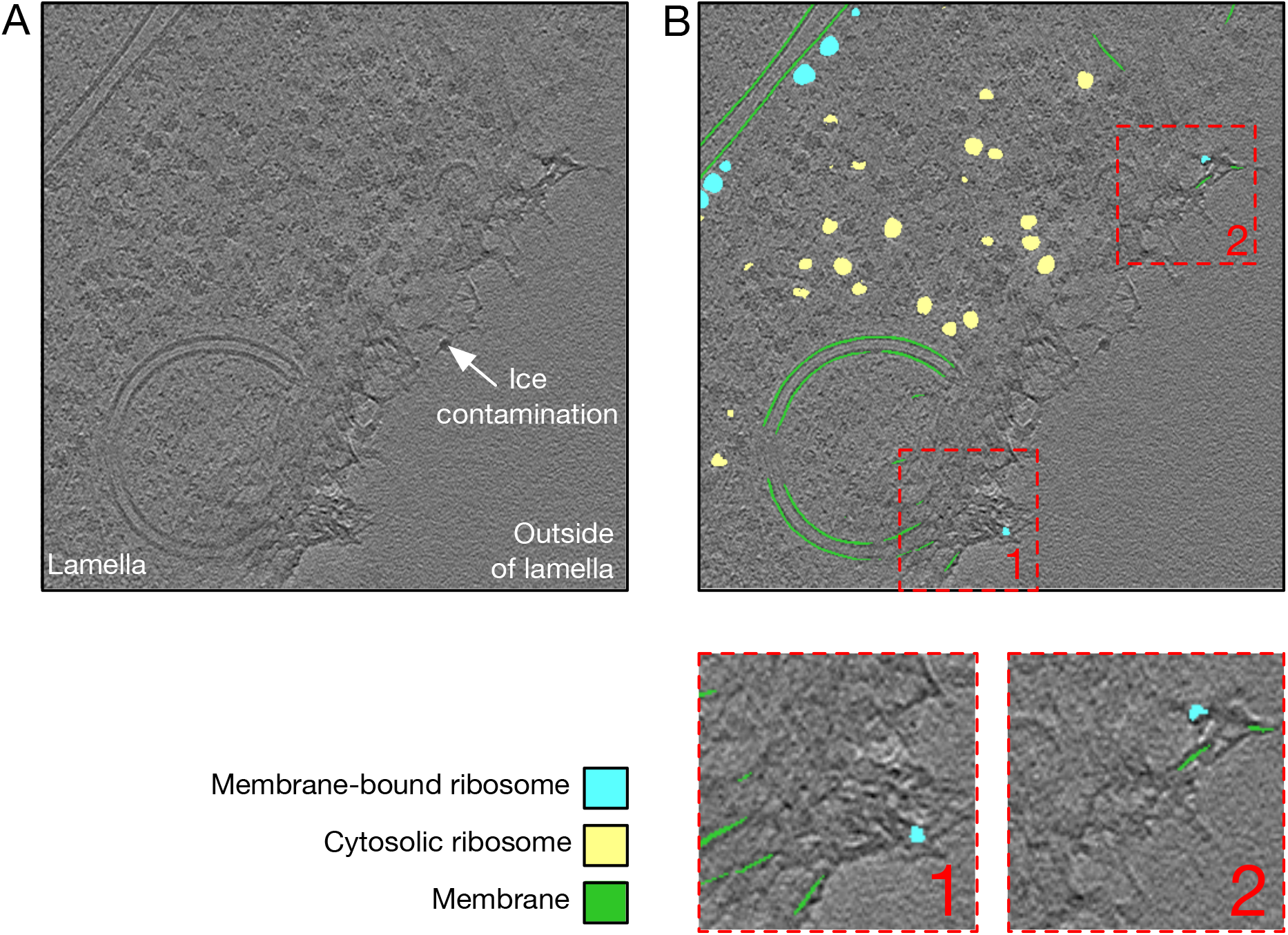
DeepFinder handles ice contamination on the lamella surface. (A) Tomogram slice depicting the border of a FIB-milled lamella. The lamella contains a *Chlamydomonas reinhardtii* cell, with a lamella border suffering from ice contamination. (B) Tomogram slice with superimposed DeepFinder segmentation. Most of the ice contamination artifacts have been correctly classified as “background”. Nonetheless, some miss-classifications exist, as can be observed in the zoomed-in boxes (in dashed red). In boxes 1 and 2, DeepFinder confuses some artifacts with membranes, and some features are wrongly classified as membrane-bound ribosomes. Such miss-classifications can be filtered out, either by masking out the outside of the lamella, or by ejecting segmented object that are too small (using the “cluster size” attribute given by the clustering step of the DeepFinder analysis stage).

**Figure 10:**
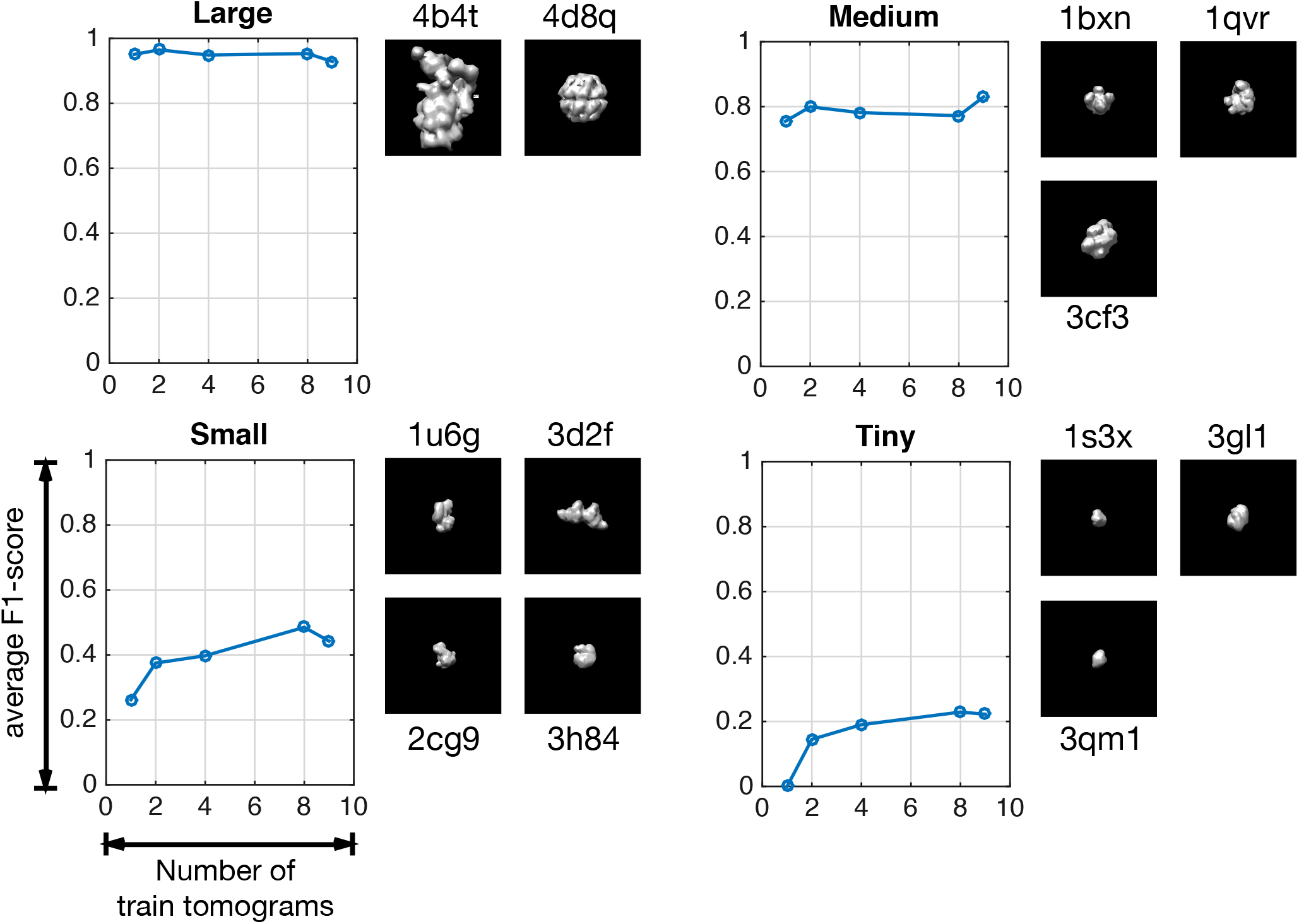
Evolution of *F*_1_-scores with respect to sizes of the training sets (number of tomograms) on the synthetic *SHREC’19* dataset (12 classes). The purpose of this figure is to give an estimation of the amount of annotated data needed to identify macromolecules. This amount depends on the size of the target macromolecule: the smaller the target, the more annotations are required. Each tomogram contains in average 208 macromolecules per class. The macromolecules have been categorized into 4 groups (large, medium, small and tiny).

**Figure 11:**
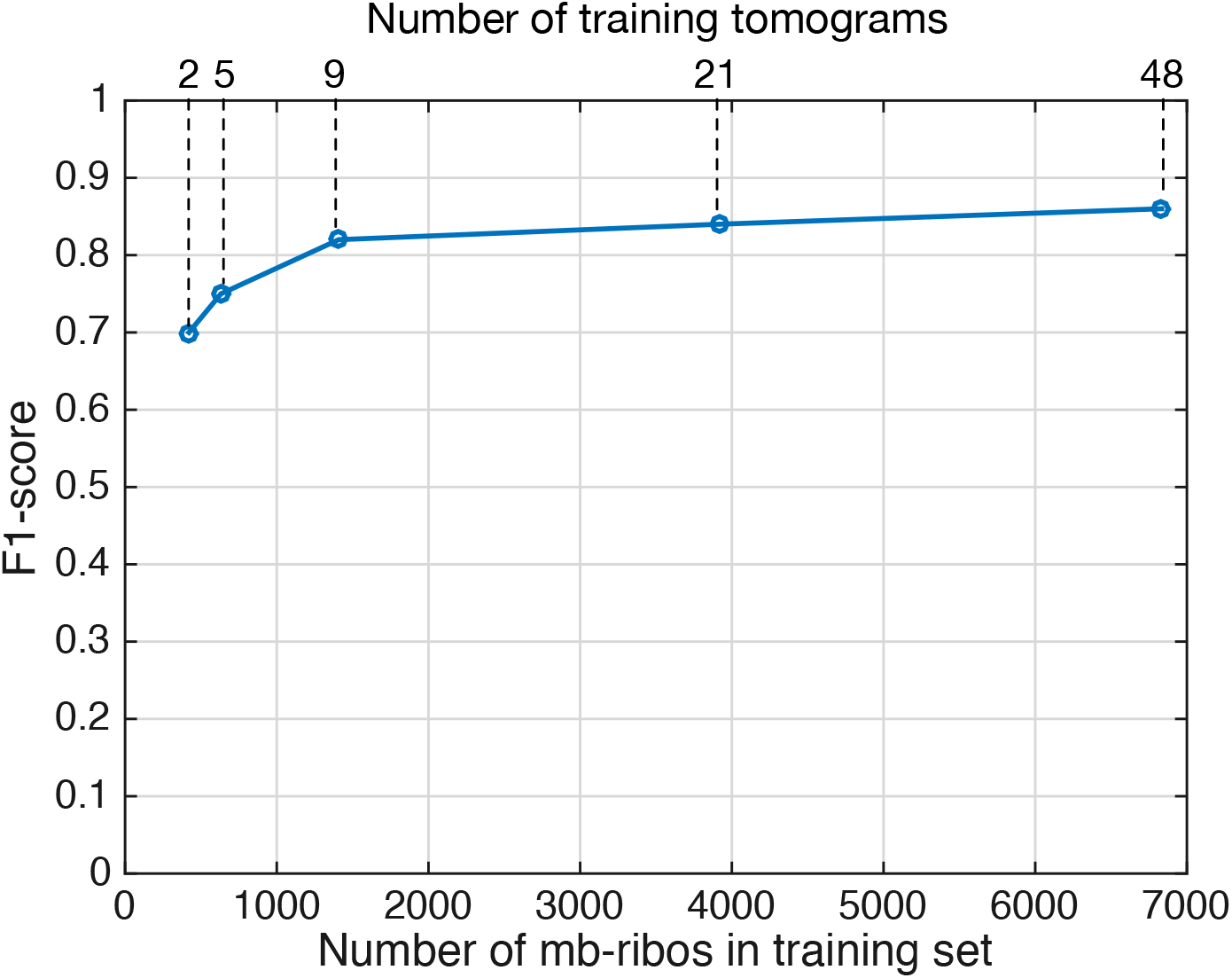
Evolution of *F*_1_-score with respect to the sizes of the training sets on real cryo-ET dataset #2, *Chlamydomonas reinhardtii* (3 classes). The *F*_1_-scores are obtained by comparing the membrane-bound ribosomes (*mb-ribo* particles) found by DeepFinder to expert annotations. As shown in Figure 10, this figure is intended to provide an estimate of the quantity of training data required to achieve a competitive result. It appears that this quantity is 1400 ribosomes (9 tomograms), which is a typical size for a cryo-ET dataset. On first glance, this estimate seems to contradict the estimates in Figure 12: the numbers do not coincide (the curve labeled “Large” estimates that quantity at 208 particles). Note that *SHREC’19* is a synthetic dataset, composed of 12 classes. Here, we are dealing with a real dataset consisting of 3 classes (membrane, membrane-bound ribosome and cytosolic ribosome). It appears that having a larger number of classes enables the use of smaller learning sets. On the other hand, the case of real data is more difficult, notably because of the presence of “label noise” (errors due to the annotation pipeline).

**Figure 12:**
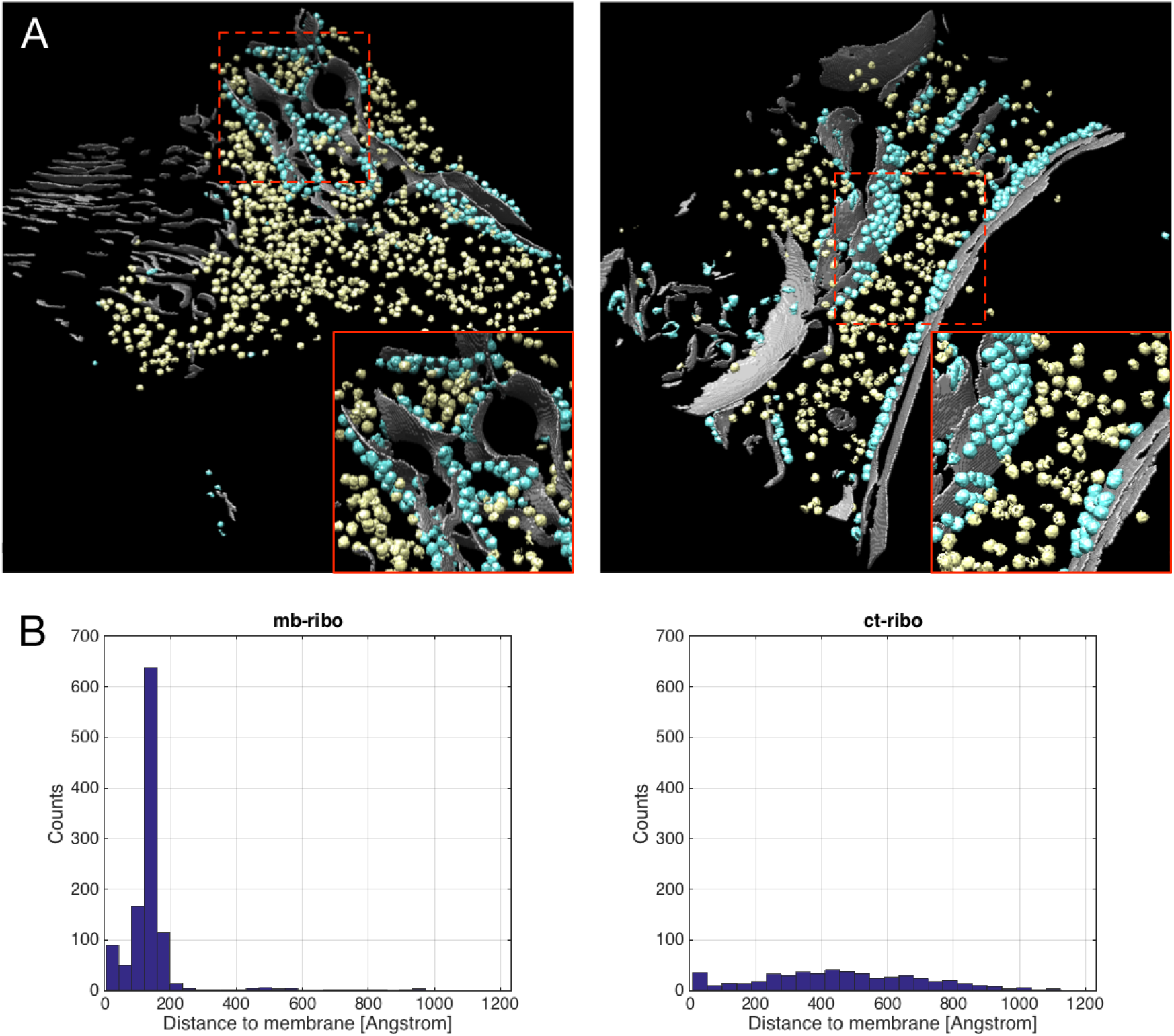
Analysis of localization of *mb-ribo* and *ct-ribo* particles in a *Chlamydomonas reinhardtii* cell (Dataset #2). (A) 3D voxel-wise classification of experimental tomograms obtained by DeepFinder. Visually, the classification makes sense: most identified membrane-bound ribosomes (in blue) are close to the cell membranes (in gray), while the cytosolic ribosomes are not. (B) Histograms of distances for *mb-ribo* particles (left) and *ct-ribo* particles (right) with respect to the nearest cell membrane computed from the test set. The histogram mode maximum for *mb-ribo* particles is located at 136.8Å, which corresponds to the ribosome radius.

**Figure 13:**
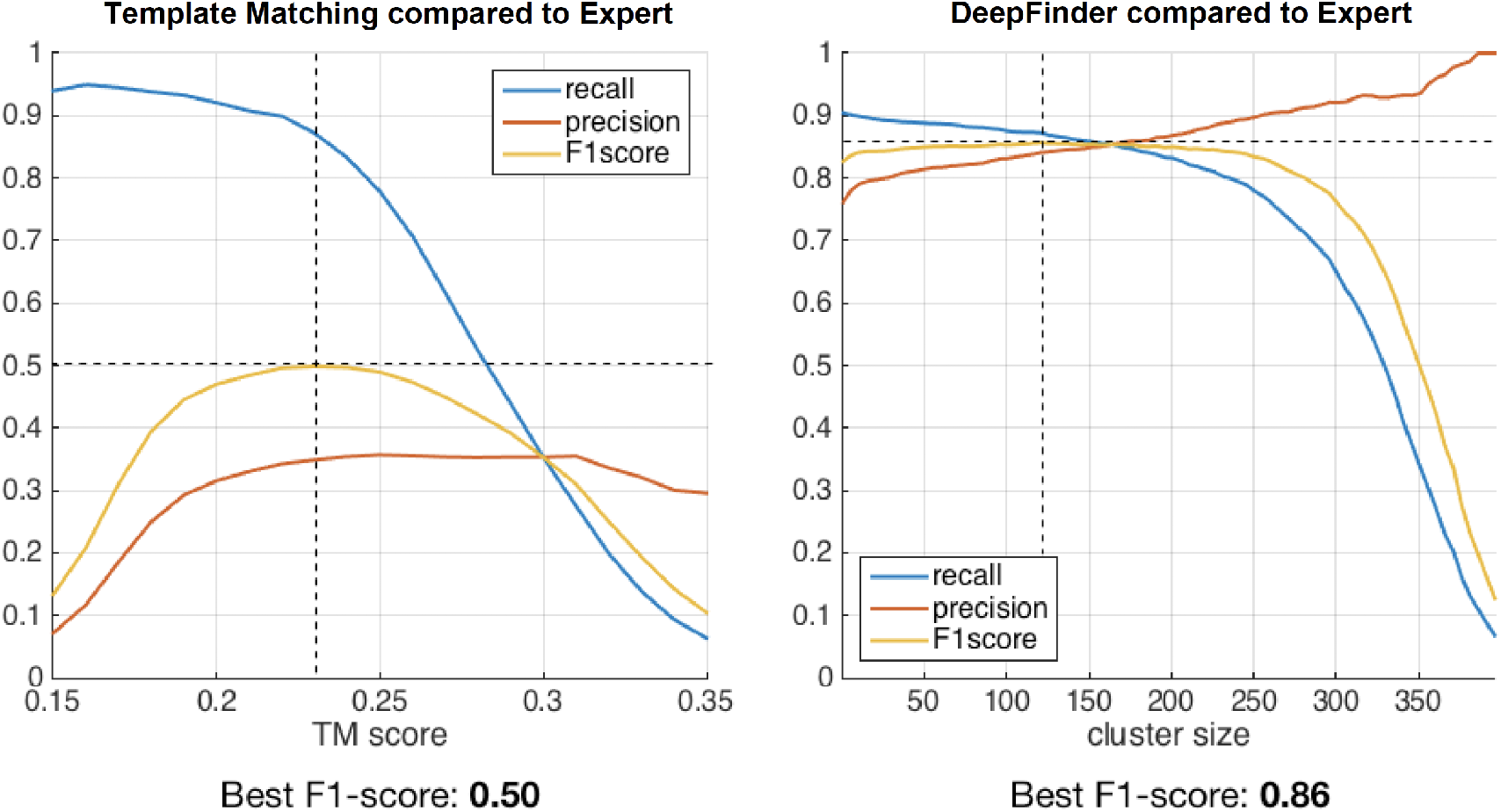
Quantitative analysis of overlap with expert annotations (Dataset #2). We varied the thresholds of template matching (left) and DeepFinder (right) to compute the *Recall, Precision* and F1-score curves. The threshold parameter for template matching is the constrained correlation coefficient, and for DeepFinder is the cluster size, which corresponds to the macromolecule radius (in voxels). Template matching and DeepFinder have good *Recall* values but template matching has a lower *Precision* than DeepFinder. This suggests that template matching can be recommended to select many candidates, but a time-consuming post-classification is required to improve *Precision*. DeepFinder has much higher *Precision* values, which confirms the results from the synthetic dataset (SHREC’19 challenge).

**Figure 14:**
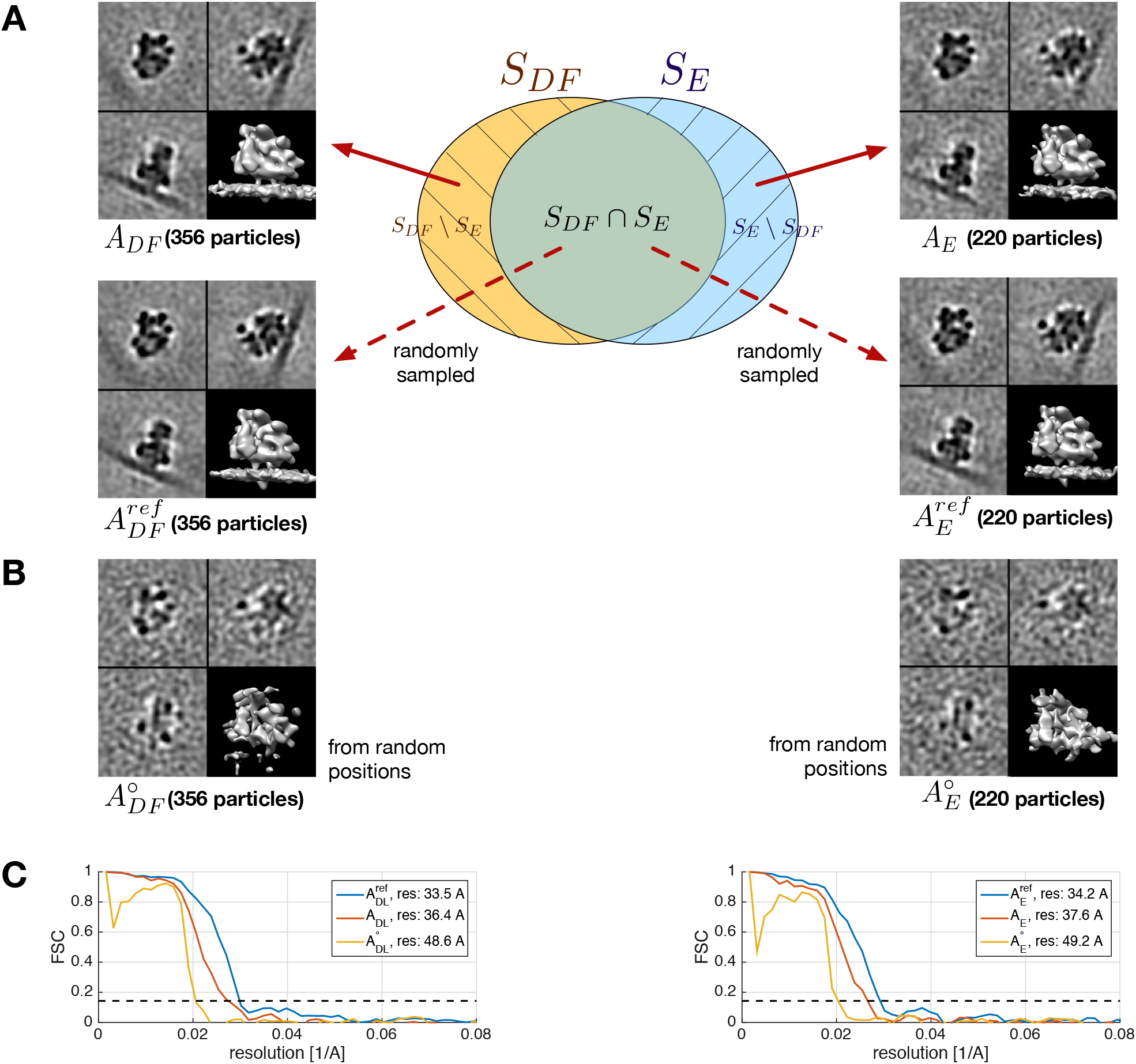
Analysis of consensus decisions and overlap sets (Dataset #2). (A) The central diagram represents the overlap between the *mb-ribo* sets *S_DF_* (found by DeepFinder) and *S_E_* (annotated by expert). Thus, *S_E_* ⋂ *S_DF_* is the subset of *mb-ribo* particles found by both DeepFinder and the experts, *S_DF_* \ *S_E_* is the subset of *mb-ribos* found by DeepFinder only (and missed by the experts), and *S_E_* \ *S_DF_* is the subset of *mb-ribo* particles found by the experts only (and missed by DeepFinder). The origin of red arrows pointing to the subtomogram averages 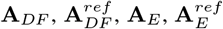 indicate the particles subsets that served to compute the averages. A ribosome density is clearly visible in **A**_*DF*_, therefore one can safely assume that the FP rate in *S_DF_* \ *S_E_* is low. (B) The subtomogram averages 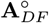 and 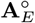 have been computed using subtomograms sampled from random positions. These averages serve to estimate a lower bound for the FSC curve. The correlation values equal or below this bound are considered “noise” values, and are caused by alignment bias^52^. (C) The averages 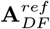 and 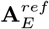 have both led to a higher resolution than **A**_*DF*_ and **A**_*E*_, implying that the *mb-ribo* particles in the set *S_DF_* \ *S_E_* and in the set *S_E_* \ *S_DF_* are more heterogenous than the *mb-ribo* particles in the set *S_DF_* ⋂ *S_E_*. Also, **A**_*DF*_ and **A**_*E*_ have led to a higher resolution than 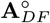 and 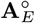, meaning that the impact of alignment bias is not significant.

**Figure 15:**
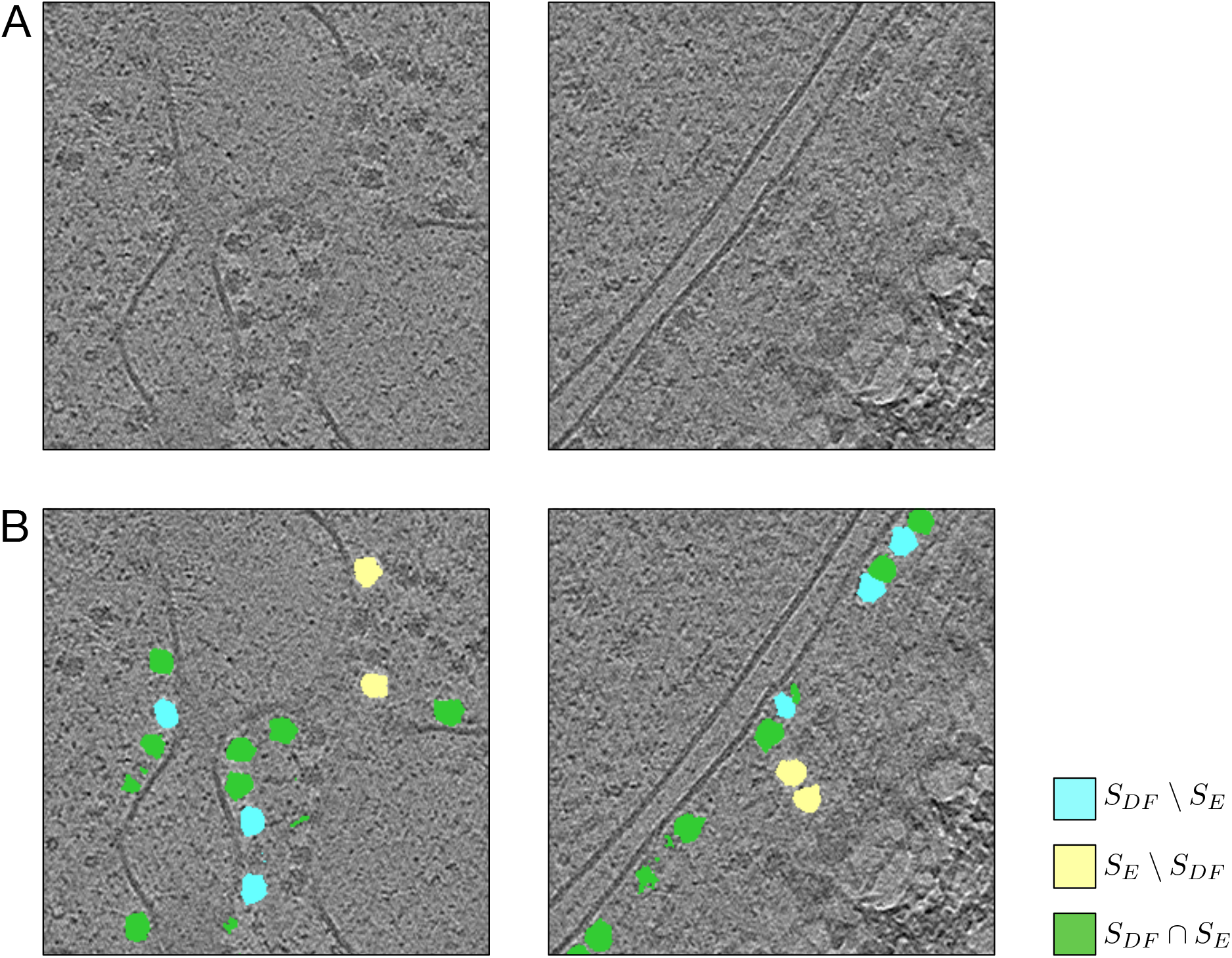
Analysis of localization consensus between DeepFinder and experts. (A) Two tomogram slice ROIs depicting *Chlamydomonas reinhardtii* cells. (B) Membrane-bound ribosomes mapped into the ROIs. In blue, the ribosomes found by DeepFinder but missed by the experts (*S_DF_* \ *S_E_*). In yellow, the ribosomes found by the experts but missed by DeepFinder (*S_E_* \ *S_DF_*). The ribosomes found by both DeepFinder and the experts (*S_DF_* ⋂ *S_E_*) are identified in green. As expected, members of *S_DF_* ⋂ *S_E_* constitute the majority of identified ribosomes. Members of *S_DF_* \ *S_E_* have a tendency of being at locations where the membrane has less contrast (B, left) or where neighboring ribosomes are close (B, right). Members of *S_E_* \ *S_DF_*, which were obtained with the expert pipeline (template matching and CPCA clustering), may also be located at positions where membrane contrast is low (B, left). Nevertheless, it appears that this pipeline has a tendency of confusing membrane-bound and cytosolic ribosomes. The proximity of ice-contamination (B, right) also seems to be a factor responsible for miss-classifications.

**Figure 16:**
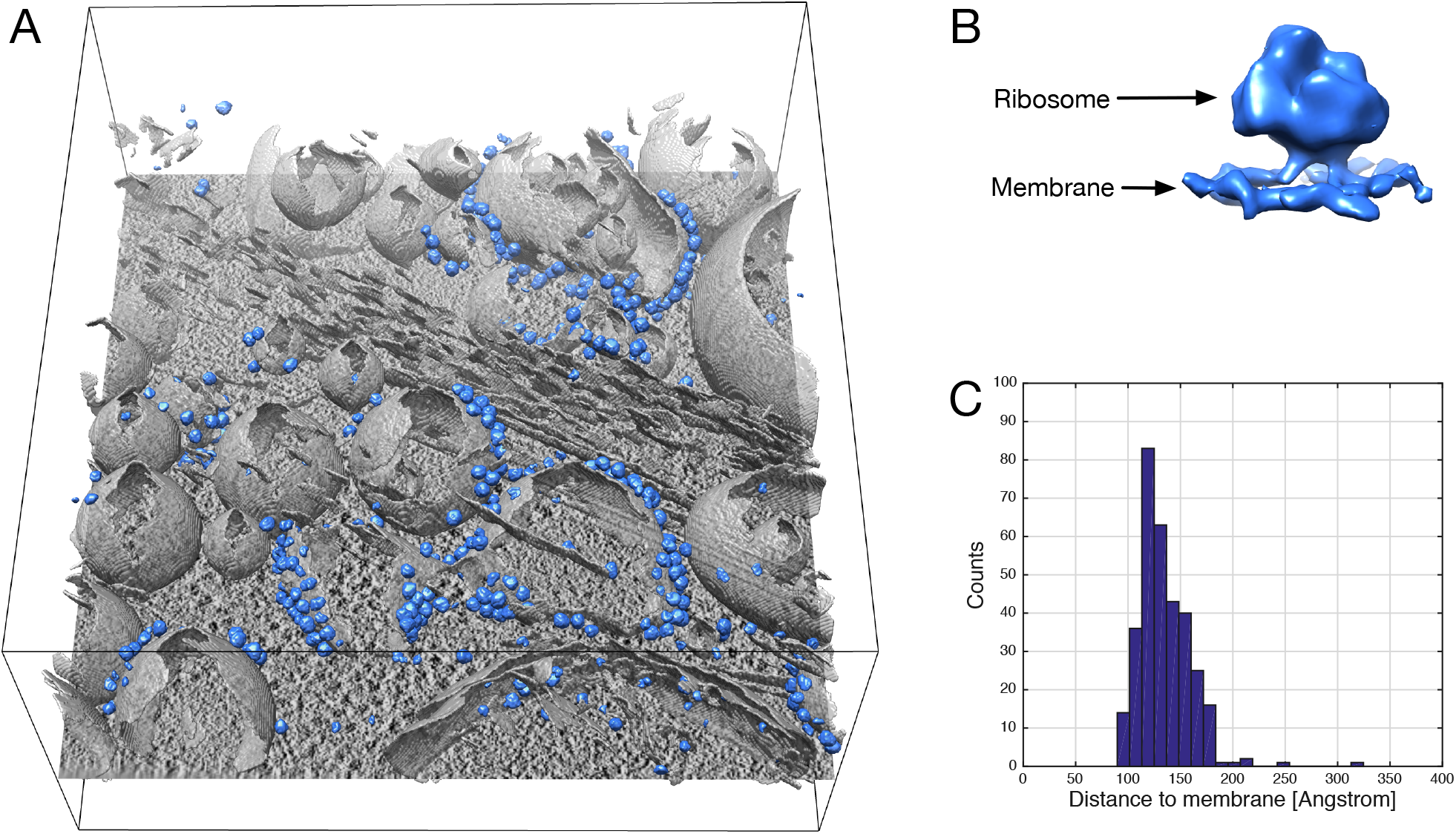
Transfer learning on P19 cells. DeepFinder has been trained on the Chlamydomonas (algae) dataset and applied on a tomogram (EMD-10439, https://www.ebi.ac.uk/pdbe/entry/emdb/EMD-10439) of P19 cells (mouse). Although the ribosome has a different structure for the two species for a given voxel size (13.68Å), the structures are similar enough for DeepFinder to identify and localize *mb-ribo* particles in a P19 cell. (A) Tomographic slice with both the superimposed segmented cell membrane (gray) and *mb-ribo* particles (blue). (B) Average density of *mb-ribo* particles. (C) Histograms of distances of *mb-ribo* particles with respect to the nearest cell membrane. The histogram mode maximum is located at 136.8Å, which corresponds to the ribosome radius.

**Figure 17:**
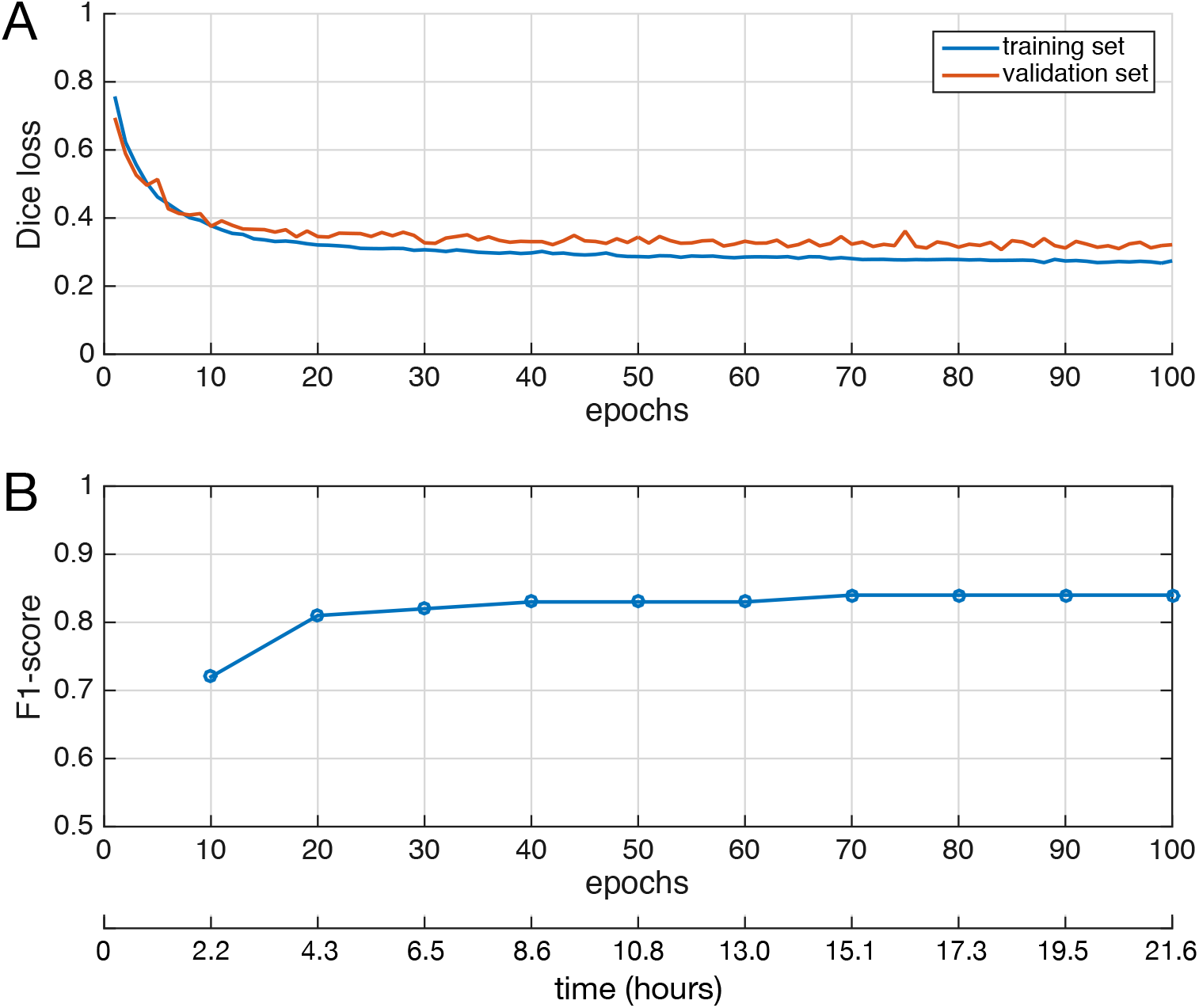
Evolution of loss and *F*_1_-score with respect to training iterations (Dataset #2). (A) The loss, which quantifies the segmentation quality, is computed for the training set, as well as for the validation set. Comparing both curves allows assessment of the generalization capabilities of our 3D CNN method. The curves for both sets should ideally overlap, otherwise it indicates overfitting (the network memorizes trained samples instead of learning discriminating features). One epoch equals to 100 training iterations. (B) The Fi-score, which quantifies the localization performance, computed on the test set. The time axis has been obtained using a Tesla K80 GPU. The curve indicates that competitive particle picking results are obtained after 20 epochs, or 4.3 hours with the required GPU.

## Appendix A

### Description of Template Matching method

We provide herein a short description of the Template Matching method (see PyTom implementation: http://strubi.chem.uu.nl/pytom/doc/pytom/tutorial.html) used in our experiments and SHREC’20 challenge^34^, and explained in detail in (Hrabe et al., 2012).

Template matching correlates a structural model of the target molecule (template) with a tomogram. The voxels corresponding to high cross-correlation coefficients are potential locations of the molecule of interest. In (Hrabe et al., 2012), the authors utilize the Local Constrained Cross Correlation (LCCC) for localizing candidate positions and apply Fourier Transform for fast computation. The computation of LCCC involves a binary mask or a weighting mask. In PyTom, non-symmetrical masks are supported and the Euler angle space is uniformly sampled to improve computational efficiency.

As described in the SHREC’20 challenge^34^, the templates were modulated in the frequency domain using a standard CTF curve at 3 *μ*m defocus. Frequencies beyond the first CTF-zero were set to 0. Spherical template masks with Gaussian smoothed edges based on the thresholded electron density were used for normalization for the cross-correlation value. We selected the best candidates with the highest cross-correlation score for each class. Then, overlapping candidates were filtered out. Two particles were considered as overlapping if the distance between their centers was smaller than the sum of their radii.

